# Suppressing chemoresistance in lung cancer via dynamic phenotypic switching and intermittent therapy

**DOI:** 10.1101/2020.04.06.028472

**Authors:** Arin Nam, Atish Mohanty, Supriyo Bhattacharya, Sourabh Kotnala, Srisairam Achuthan, Kishore Hari, Anusha Nathan, Govindan Rangarajan, Erminia Massarelli, Herbert Levine, Mohit Kumar Jolly, Prakash Kulkarni, Ravi Salgia

**Affiliations:** Department of Medical Oncology and Therapeutics Research, City of Hope National Medical Center, Duarte, CA 91010, USA; Translational Bioinformatics, Center for Informatics, Department of Computational and Quantitative Medicine, City of Hope National Medical Center, Duarte, CA 91010, USA; Center for Informatics, Division of Research Informatics, City of Hope National Medical Center, Duarte, CA 91010, USA; Center for BioSystems Science and Engineering, Indian Institute of Science, Bangalore 560012, India; Department of Mathematics, Indian Institute of Science, Bangalore 560012, India; Center for Neuroscience, Indian Institute of Science, Bangalore 560012, India; Department of Physics, Northeastern University, Boston, MA 02115, USA; Department of Bioengineering, Northeastern University, Boston, MA 02115, USA

**Keywords:** chemoresistance, cisplatin, lung cancer, evolutionary game theory, group behaviour, persister trait, phenotypic switching

## Abstract

A major challenge in cancer therapy is drug resistance, which is typically attributed to acquired mutations and tumor heterogeneity. However, emerging evidence suggests that dynamic cellular interactions and group behavior also contribute to drug resistance, although, the details of such mechanisms are poorly understood. Here, by combining real time cellular growth data with mathematical modeling, we showed that the cisplatin-sensitive and tolerant lung cancer cells when co-cultured in cisplatin-free and cisplatin-treated environments, exhibit drastically different group strategies in response to environmental changes. While tolerant cells exhibited a persister-like behaviour and were attenuated by sensitive cells, sensitive cells ‘learned’ to evade chemotherapy from tolerant cells when co-cultured. Further, tolerant cells could switch phenotypes to become sensitive, although high cisplatin concentrations suppressed this switching. Finally, switching cisplatin administration from continuous to intermittent suppressed the emergence of tolerant cells, suggesting that intermittent rather than continuous chemotherapy may result in better outcomes in lung cancer.

---

Resistance to chemotherapy is a major impediment in treating cancer. Resistance is generally held to primarily arise through random genetic mutations and the subsequent expansion of mutant clones via Darwinian selection (Salgia and Kulkarni, 2018; Álvarez-Arenas et al, 2019; Greene et al, 2019). Hence, the phenomenon has been approached from a reductionist, gene-centric perspective. However, emerging evidence suggests a major role of non-genetic mechanisms in drug resistance by enabling the cancer cells to be phenotypically plastic (Dawson and Kouzarides, 2012; Jones et al, 2016; Bell and Gilan, 2020), thus increasing their adaptability and cooperation under stressful conditions (Wu et al, 2014; Kaznatcheev et al, 2019; Stanková K, 2019). Nonetheless, such mechanisms have not been fully explored and are rarely integrated into clinical trials or in precision oncology initiatives (Mirnezami et al, 2012; Cunningham et al, 2018; Stanková et al, 2019). Combining conventional therapies with treatment strategies based on cancer ecology could potentially delay or even prevent drug tolerance and eventually, drug resistance (Zhang et al, 2017; Salgia and Kulkarni, 2018; Stanková et al, 2019).

Here, we have used human non-small cell lung cancer (NSCLC) cells that are either naturally sensitive or tolerant to cisplatin as a paradigm to determine how group behaviour (competition and cooperation) may help them evade the effects of chemotherapy. Cells were cultured either by themselves (monotypic) or after they were mixed together (heterotypic) in different ratios to discern cell-autonomous and non-cell-autonomous fitness effects. Cell-autonomous fitness effects are inherent to the cell, independent of the presence of other cellular phenotypes in the ecosystem. Hence, the growth rates from monotypic cultures could potentially provide the necessary information for determining these effects (Kaznatcheev et al, 2019). Non-cell-autonomous effects on the other hand are those that allow fitness to depend on a cell’s microenvironment, including the frequency of other cell types (Marusyk et al, 2014). Therefore, growth rates need to be measured in competitive fitness assays over a range of seeding ratios. The two cell populations were monitored in real time, and competition and cooperation were mathematically modeled applying a new kinetics-based stress-mediated phenotypic switching model (PSMSR) with a game theoretic underpinning, and estimating the parameters through the best fit of measurements obtained from the experiments.

## Results

### Cisplatin-sensitive and tolerant cells demonstrate different behaviours in monotypic and heterotypic cultures

Cisplatin-sensitive H23 and cisplatin-tolerant H2009 NSCLC cell lines permanently tagged with red fluorescent protein (RFP) and green fluorescent protein (GFP), respectively (Mohanty et al, 2019), were co-cultured in 10 cm dishes and monitored in real time (**Supplementary Fig. 1A**). To discern differences in their behavior, the two cell cultures were grown separately (monotypic) or by mixing them in a 1:1 ratio (heterotypic) 12 h before the start of the experiment. Alternatively, they were mixed in a 1:1 ratio and co-cultured for 3 weeks before determining their growth rates (**Fig. 1A, schematic**). There was no significant difference in the growth rates of the sensitive cells in any of the conditions tested, and they were equally sensitive to 5 μM cisplatin in all three conditions (**Fig. 1A, left panel in red**). Similarly, when tolerant cells were cultured by themselves or were mixed just before the start of the experiment, they did not show any significant change in growth rate or drug tolerance. However, when co-cultured for three weeks prior to the experiment, the tolerant cells showed marked reduction in cell proliferation. When they were cultured in presence of cisplatin, they showed less sensitivity compared to their counterparts cultured separately or mixed 12 h before the experiment was started. Thus, the tolerant cells appeared to exhibit a persister-like trait in absence of cisplatin but reverted to active proliferation in the presence of the drug (**Fig. 1A, right panel in green**).

**Fig. 1:**
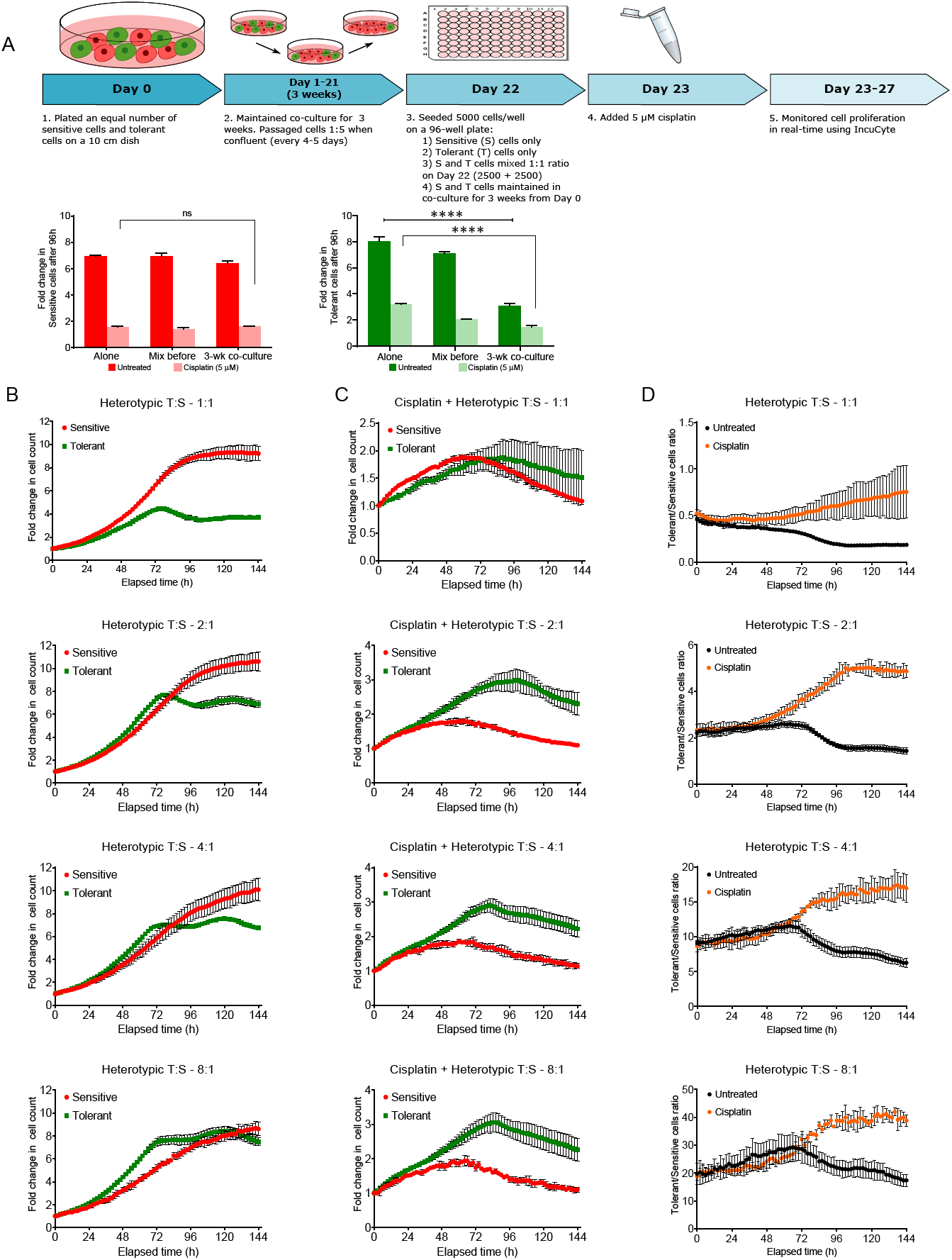
Behaviour of cisplatin-sensitive (S) and tolerant (T) NSCLC cells in 2D co-culture. (**A**) Schematic representation of the experimental design of co-culturing S and T cells in a ratio of 1:1 and collection of data points. Proliferation of sensitive (red) and tolerant (green) cells under different culture conditions in the absence or presence of cisplatin. Two-way ANOVA test (multiple comparison) showing statistical significance ****p<0.0001. (**B**) Sensitive and tolerant cells were plated in increasing T:S ratios and cultured for 3 weeks. Proliferation rate of sensitive cells (red) and tolerant cells (green) in heterotypic culture over the course of 144 hours. (**C**) Proliferation rate of sensitive cells (red) and tolerant cells (green) in heterotypic culture treated with 5 μM cisplatin over the course of 144 hours. (**D**) Change in tolerant/sensitive cells ratio with (orange) and without (black) 5 μM cisplatin over the course of 144 hours. Statistical significance information can be found in **Supplementary Table 2**.

Next, we seeded the two cell populations at different ratios and cultured them for 3 weeks. The cell number was then counted, 5000 cells were seeded per well in a 96-well plate, and the growth rates in the presence or absence of cisplatin were determined **(Supplementary Fig. 1B, schematic**). At a seeding ratio of 1:1, sensitive cells clearly suppressed tolerant cell proliferation (**Fig. 1B, upper left panel**). However, when the ratio of tolerant to sensitive cells was increased to 2:1, the proliferation rates for both cell types were almost similar for 72 h but showed a decline thereafter (**Fig. 1B, second plot from top, left panel).** When the ratio was increased further to 4:1 or 8:1, the tolerant cells proliferated at a much faster rate compared to sensitive cells, and the suppressive effect of sensitive cells was minimal (**Fig. 1B lower left bottom two panels**). However, the inhibitory effect was negligible in the presence of cisplatin. In fact, tolerant cells proliferated better in the presence of cisplatin, and increasing the seeding ratio of tolerant cells in the population also favored their growth (**Fig. 1C**).

Next, we calculated the change in the ratio of tolerant versus sensitive cells over 144 h under untreated (black line) and cisplatin-treated (orange line) conditions. We observed that, under both conditions, there was minimal change in the ratio up to 72 h and thereafter, there was a drop in the ratio for the untreated culture, indicating a decline in tolerant cell population. In contrast, there was an increase in the ratio in presence of cisplatin, suggesting a reduction in the sensitive cell population. Together, these observations suggest that after 72 h of co-culture, there was a transition in the cell behavior in the untreated condition favoring sensitive cell growth but suppressing tolerant cell growth either because of the depletion in space and nutrients or secretion of inhibitory factors (**Fig. 1D**).

To distinguish the two possibilities, we repeated the above experiment by mixing the sensitive and tolerant cells, but instead of 3 weeks we incubated the co-culture for only 12 h prior to starting the experiment (**Supplementary Fig. 2A, schematic**). We observed that sensitive cells suppressed tolerant cells at 1:1 ratio, and the suppressive effect became negligible with increase in tolerant cell ratios from 1:1 to 8:1 (**Supplementary Fig. 2B and D, and Supplementary Table 1**). Further, the tolerant cells remained dominant in the presence of cisplatin (**Supplementary Fig. 2C and D**). As the tolerant population increased, the reduction observed at 1:1 ratio appeared to favor the tolerant cells, again suggesting that tolerant cells are efficient in sensing the presence of their subtype (an apparent kin selection-like behavior). Alternatively, it is possible that, the increase in the tolerant cell numbers, diluted the inhibitory effect of a factor secreted by the sensitive cells. Therefore, we performed the experiment as above but with the cells mixed in increasing ratio of sensitive versus tolerant cells from 1:1 to 8:1 and co-cultured for 3 weeks (**Supplementary Fig. 3A, schematic**). Consistent with the previous experiments, in the absence of cisplatin, the sensitive cells suppressed the growth of tolerant cells, but in the presence of cisplatin, the tolerant cells dominated the population overall (**Supplementary Fig. 3B-D**). Together, these experiments indicated that in a heterogeneous population, dynamic competition and cooperation exists between the sensitive and the tolerant cells, and the sensitive cells dominate over tolerant cells. In contrast, the presence of cisplatin favors the survival and proliferation of tolerant cells by reducing the competition between the tolerant and sensitive cells.

### A secretory factor from sensitive cells rather than direct interaction impedes tolerant cell growth

To discern whether a physical interaction between the two cell types is necessary or sensitive cells secreted an ‘inhibitory factor’ to attenuate tolerant cell growth, the cells were grown in conditioned medium from sensitive or tolerant cells instead of the fresh cell culture medium. Conditioned medium from the sensitive cells impeded proliferation of tolerant cells by ∼4.5-fold. In contrast, conditioned medium from tolerant cells had no significant effect on the growth of sensitive cells, alluding to the presence of an inhibitory factor in the conditioned medium from the sensitive cells (**Fig. 2A**). To determine whether the putative inhibitory factor acted in a dose-dependent fashion, we seeded the two cell types at varying densities (2,500-20,000 cells/well) and compared the fold change of inhibition in the growth of tolerant cells grown in conditioned medium from sensitive cells compared to the cells grown in fresh medium. As expected, at a lower seeding density, the conditioned medium had a greater inhibitory effect (approximately 6-fold) on the tolerant cells and reduced in a dose-dependent fashion with an increase in the number of tolerant cells (**Fig. 2B**).

**Fig. 2:**
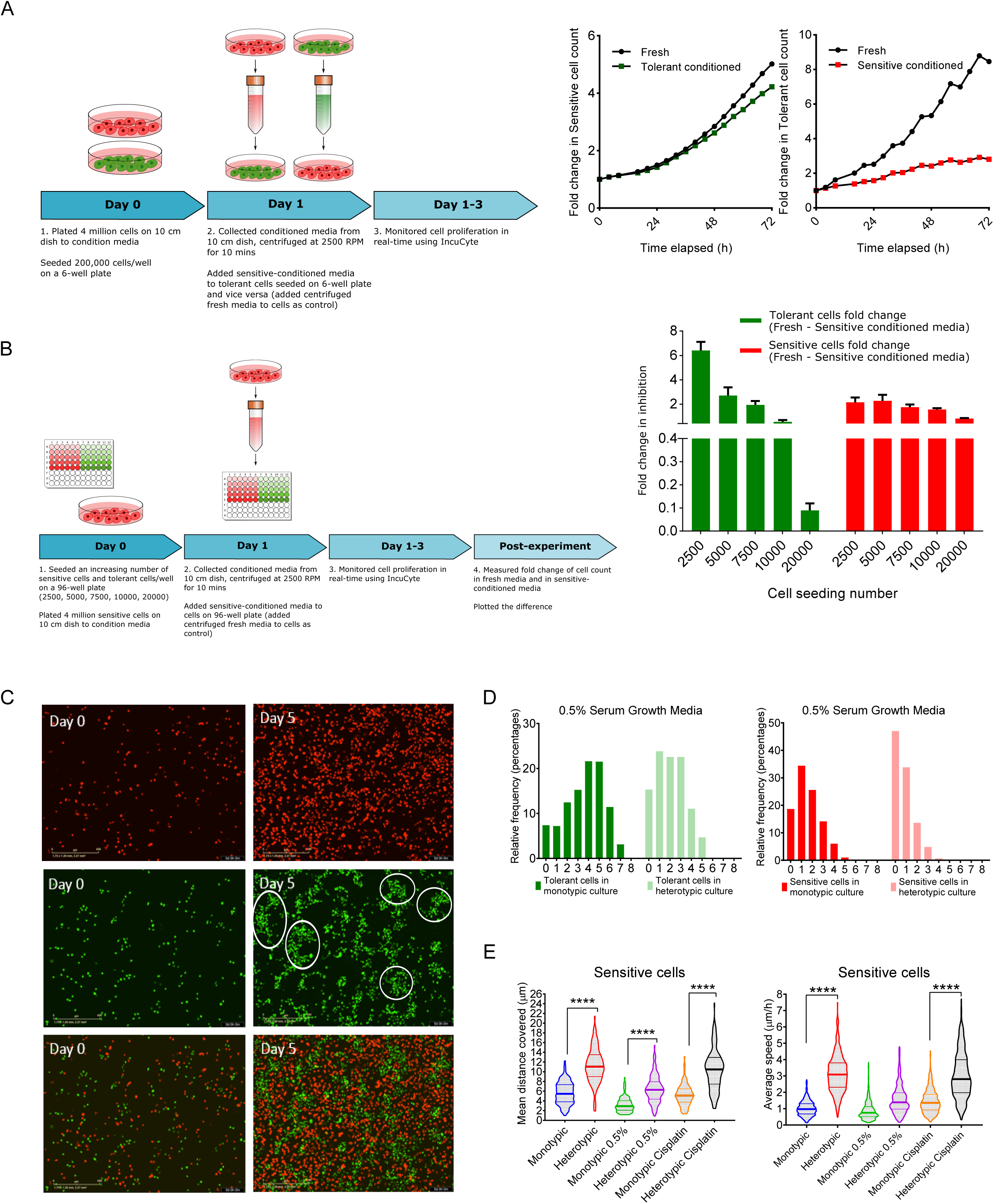
Transition of behaviors in cisplatin-sensitive and tolerant cells in 2D co-culture. (**A**) Effect of conditioned medium on growth of cisplatin-sensitive and cisplatin-tolerant cells compared to cells cultured in fresh medium. Schematic representation of the experimental design to understand the effect of conditioned medium from sensitive cells on tolerant cells and vice-versa and analyzed data (graphs on the right). (**B**) Effect of conditioned medium of sensitive cells on the growth of cisplatin-sensitive and cisplatin-tolerant cells seeded at different densities. Schematic representation of the protocol and the analyzed data (bar diagrams on the right). (**C**) Differences in the spatial behavior patterns of the cisplatin-sensitive and cisplatin-tolerant NSCLC cells in 2D culture conditions. The tolerant cells exhibited group behavior (highlighted with white circles). (**D**) Group behavior among tolerant cells is enhanced under stress. The Y axis depicts the relative frequency of the nearest neighbor when sensitive and tolerant cells were grown in monotypic or heterotypic culture under low 0.5% serum. (**E**) Effect of co-culture on sensitive cells speed and distance. Violin plots indicate the mean distance covered and the relative speed of the cisplatin-sensitive cells under various culture conditions. Statistical significance information can be found in **Supplementary Table 3**.

### Sensitive and tolerant cells exhibit different spatial patterns in the absence and presence of each other

We plated sensitive and tolerant cells separately in 10 cm dishes to track their behaviour over a period of 5 days. In the absence of cisplatin, tolerant cells tended to grow in close proximity (groups), whereas the sensitive cells appeared more dispersed (**Fig. 2C**), suggesting that the tolerant cells were able to communicate and cooperate to form more drug-tolerant colonies that proliferate rapidly (**Fig. 2C**). Even in the presence of cisplatin, the tolerant cells showed cooperation (tended to form clusters), while the sensitive cells did not (**Fig. 2D**).

To measure clustering behavior, we used the QuPath software and identified the number of nearest neighbors across each cell under various conditions (**Supplementary Fig. 4A**). As the clustering increased, the number of nearest neighbors also increased. To rule out the possibility that the increase may be due to cell confluence, we cultured the cells in serum-deprived condition where proliferation is attenuated. Regardless, the clustering behavior of tolerant cells increased as the fraction of cells having nearest neighbor >3 increased (**Fig. 2D, left panel dark green bar graph**). However, clustering was perturbed when sensitive cells were co-cultured with tolerant cells, which may explain an increase in the fractions of cells having fewer nearest neighbors ≤ 3 (**Fig. 2D, left panel light green bar graph**). Similarly, we also observed the fraction of cells with 0 neighbors increased by just mixing the tolerant and sensitive cells in a 1:1 ratio and the fewer the number of nearest neighbours, the weaker was the proliferation rate of the resistant cells (**Supplementary Fig. 4B, green graph**). In contrast, the sensitive cells were unaffected by reduction in the nearest neighbor. When cisplatin was added to the co-culture, we also observed an increase in the fraction of cells with nearest neighbor <3. Under the same condition however, the sensitive cells had the highest number of cells which have 0 neighbors (**Supplementary Fig. 4C, red graph**). Thus, having 0 neighbors appeared beneficial to sensitive cells in untreated condition as it gives them more freedom, but in stressful conditions (like cisplatin or low serum), having 0 neighbors is not beneficial.

Next, we asked how co-culturing the cells affects sensitive cell behavior. To this end, we tracked ∼200 cells by taking images every 20 minutes for 24 h and calculated the distance they covered and the mean speed **(Supplementary Fig. 5A**). Much to our surprise, in the presence of tolerant cells, the sensitive cells increased their migratory speed and mean distance covered (**Fig. 2E**). This change in their migratory behavior may indicate a synchronization of the sensitive cells in the presence of the competing tolerant cells. We then sought to relate the changes in the migratory behavior of the sensitive cells to changes in their gene expression in presence of tolerant cells. We co-cultured the sensitive and tolerant cells for 5 days then separated the sensitive and tolerant cells by FACS sorting followed by RNA extraction and sequencing. RNA expression profiles of sensitive cells cultured as monotypic or heterotypic were compared to tolerant cells cultured as monotypic culture. Using ingenuity pathway analysis software, we identified a total of 1500 genes in sensitive cells which were either directly or indirectly involved in cell movement in the heterotypic culture leading to an increase in cell movement. In contrast, only 150 genes associated with cell movement were affected in monotypic culture (**Supplementary Fig. 5B)**. Of the 1500 genes, a set of 9 common genes was present in both comparisons. The expression of these 9 genes switched from low in monotypic culture (blue heat map) to high expression in heterotypic culture (red heat map) which correlated with increased cell movement and migration **(Supplementary Fig. 5C**).

### Intermittent therapy can sustain a population of cisplatin-sensitive tumor cells while attenuating the proliferation of resistant cells

Since the proliferation of the tolerant cells was remarkably impeded when co-cultured with sensitive cells for prolonged periods (3 weeks) prior to cisplatin treatment, we asked if continuous or intermittent treatments would differentially affect a mixed population of the two cell types. Toward this end, we mixed the two cell types at 1:1, 2:1, and 4:1 (sensitive:tolerant) ratios on Day 0 and on the following day (Day 1), added a sublethal dose of cisplatin (1 μM) to the medium (**Fig. 3A, schematic**). Two days later, on Day 3, the cultures were passaged 1:5. One portion of the cells was again treated with 1 μM cisplatin and was maintained in cisplatin containing medium for the remainder of the experiment. We refer to this arm of the experiment as the ‘continuous treatment’ (**Fig. 3A, schematic**). The other portion was maintained in medium without cisplatin for 10 more days to allow the cells to proliferate. After 10 days (Day 13), this group of cells was again split into two equal portions. While one portion was maintained in medium with no cisplatin (Intermittent I cycle), the other portion was treated with 1 μM cisplatin for 4 days, washed once, and maintained in fresh medium with no cisplatin for the remainder of the experiment. We refer to this arm as Intermittent II cycle. The number of each cell type was determined in all 3 experimental sets for the course of the entire experiment (43 days) (**Fig. 3A, schematic**).

**Fig. 3:**
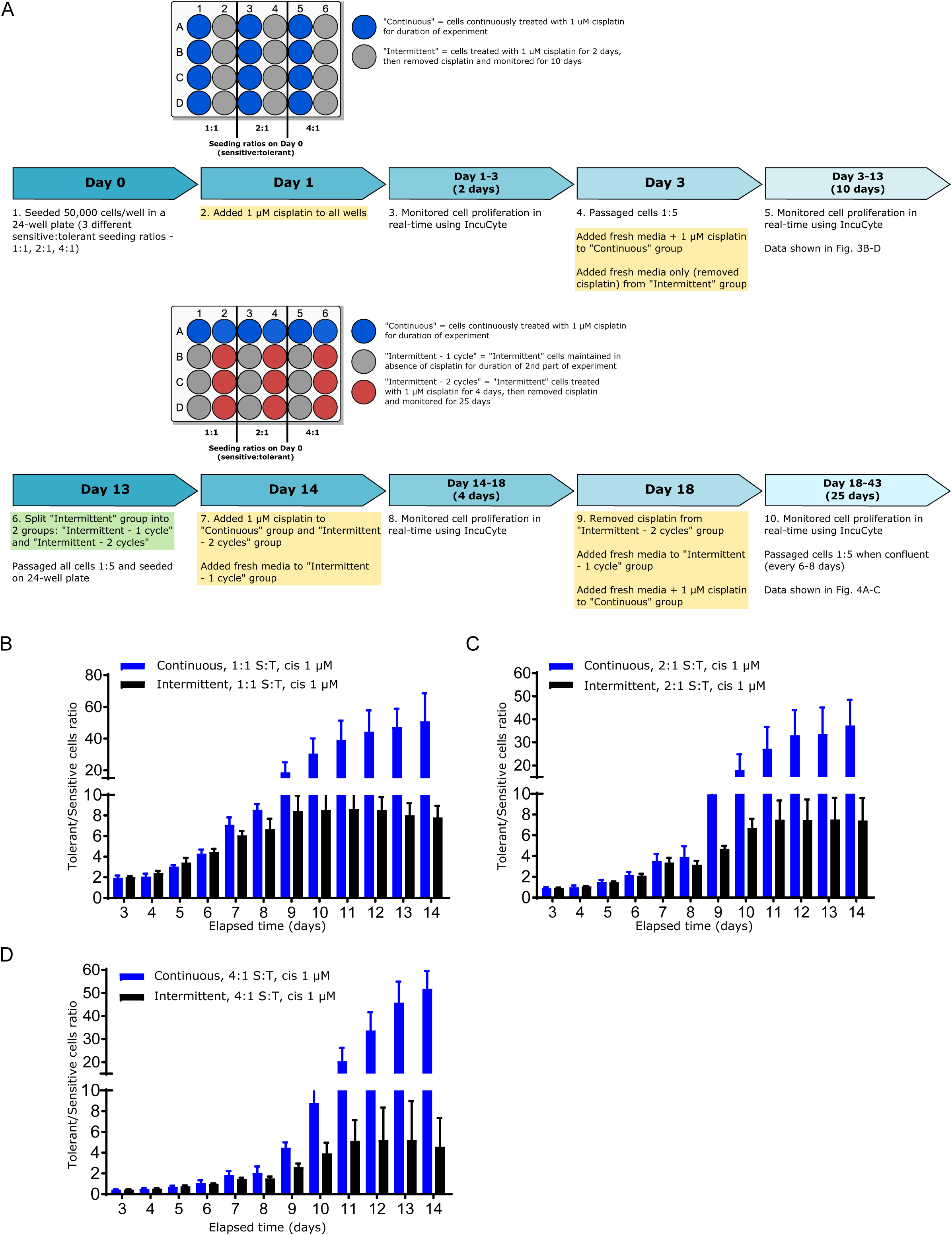
Intermittent therapy. (**A**) Schematic representation of the experimental design of continuous versus intermittent therapy in 2D cultures. Sensitive and tolerant cells were seeded in a ratio of 1:1, 2:1, and 4:1 in a 24-well plate and allowed to adhere overnight as depicted in the plate map. After 24 h (Day 1), the cells were treated with a sublethal dose (1 µM) of cisplatin for two days. After 2 days (Day 3), all cells were passaged 1:5. The “Continuous” group received fresh media containing 1 µM cisplatin. The “Intermittent” group received fresh medium only (no cisplatin). Cells were allowed to grow for 10 days and proliferation was monitored in real-time using IncuCyte. After 10 days (Day 13), cells were trypsinized. The “intermittent” group was split into 2 groups: “Intermittent – 1 cycle” and “Intermittent – 2 cycles”. All cells were passaged 1:5 and re-seeded in a 24-well plate as depicted in the plate map. After 24 h (Day 14), the “Continuous” group and “Intermittent – 2 cycles” group were treated with 1 µM cisplatin. Only fresh media was added to the “Intermittent – 1 cycle” group. Cell proliferation was monitored in real-time for 4 days using IncuCyte. After 4 days (Day 18), cisplatin was removed, and fresh media was added to “Intermittent – 2 cycles” group. Fresh media was added to “Intermittent – 1 cycle group” and fresh media containing 1 µM cisplatin was added to “Continuous” group. Cell proliferation was monitored in real-time for 25 days using the IncuCyte Live Cell Imaging System. The experiment was terminated after 25 days (Day 43). (**B-D**) Bar graph showing the ratio of tolerant versus sensitive cell population over a period of 10 days. The cell ratio for the “Continuous” group wherein the cells were continuously treated with cisplatin is shown in blue and the ratio for the “Intermittent” group wherein the cells were treated with cisplatin for 2 days and released in fresh medium (intermittent) is shown in black.

Within 10 days, we observed the ratio of tolerant to sensitive increased by 50- to 100-fold for the initial seeding density of 1:1, 2:1 and 4:1, respectively, under continuous treatment. On the other hand, the ratio for the intermittent therapy increased only 3- to 8-fold (**Fig. 3B-D**), suggesting that sensitive cells were able to recover and proliferate. We prolonged intermittent therapy by splitting the cells growing in cisplatin-free media post cisplatin treatment into two sets: ‘Intermittent 1 cycle’ and ‘Intermittent 2 cycles’ as described above. We then asked if the sensitive cells once exposed to cisplatin would overgrow the tolerant cells to recapitulate the data shown in our previous co-culture experiments. We continued the culture for 24 days, to let the cells grow and once confluent, passaged 1:5 every 6-8 days. We observed that the tolerant vs. sensitive ratio fell to approximately 2, 1.6 and 0.4 for initial seeding density of 1:1, 2:1 or 4:1, respectively, for the cells treated only once with cisplatin on Day 3 (Intermittent – 1 cycle) (**Fig. 4A-C, black line**). In contrast, the ratio of cells that received the 2^nd^ dose of cisplatin treatment (‘Intermittent – 2 cycles’) and were allowed to recover in fresh media, did not show any decrease in the tolerant population and maintained in a ratio of 100/800 (**Fig. 4A-C, red line**).

**Fig 4:**
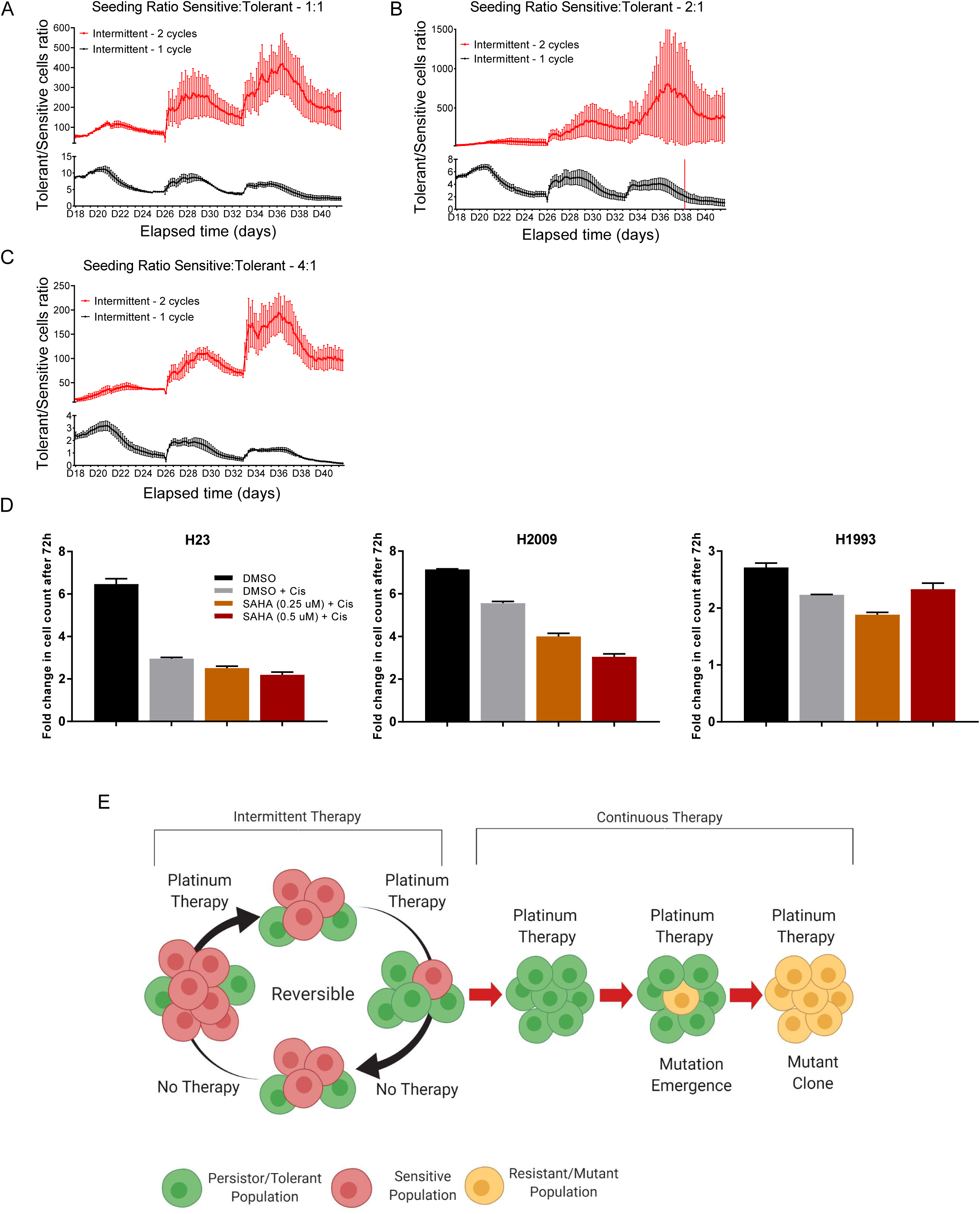
Tolerant cells reversibly switch their phenotype to become sensitive with intermittent therapy. (**A-C**) Media from “Intermittent – 2 cycles” group was removed after 4 days of cisplatin treatment and replaced with fresh medium and the cells were allowed to grow until confluent. These cells were monitored in real-time to determine the ratio of tolerant vs sensitive over the course of 25 days. Similarly, the cells that only received cisplatin once (“Intermittent – 1 cycle”) throughout the experiment were also followed for 25 days. (**D**) Effect of suberoylanilide hydroxamic acid (SAHA) on cisplatin-sensitive (H23), tolerant (H2009), and resistant (H1993) cells, demonstrating that tolerant cells can reversibly switch their phenotype to become sensitive. (**E**) A model depicting the emergence of truly cisplatin resistant cells from cisplatin tolerant cells as a result of continuous therapy. The model implies that resistance is due to irreversible mutations and their selection while tolerance is reversible and therefore, due to non-genetic mechanisms. Statistical significance information can be found in **Supplementary Table 4**.

### Epigenetic modulation can distinguish drug sensitivity, tolerance and resistance in lung cancer

Unlike in microbiology where the terms persistence, tolerance and resistance are well defined (Brauner et al, 2016), rigorous definitions of these terms are lacking in cancer (Salgia and Kulkarni, 2018). In bacteria, persisters are defined as transiently tolerant variants that allow populations to avoid eradication by antibiotics (Balaban et al, 2019). Persistence is linked to preexisting heterogeneity in the population and is modulated by phenotypic switching in normally growing cells. Thus, to determine the possibility that one can distinguish the different phenotypes insofar as response to cisplatin treatment is concerned, we selected a third NSCLC cell line H1993 based on the IC50 values for cisplatin from the Drug Sensitivity in Cancer Database (http://www.cancerrxgene.org). This database indicated the IC50 values for the H2009 and the H23 cell lines as 38 μM and 5 μM, respectively. In contrast, the IC50 for H1993 is listed as 308 μM. We reasoned that the H23 cells may be designated as sensitive, while the H1993 may represent truly resistant cells and perhaps, may be irreversible, simply because of the extremely high IC50 values. On the other hand, we conjectured that the H2009 cells may represent the tolerant cells and perhaps may be reversible to become sensitive to lower concentrations of cisplatin, much like the H23 cells.

To test these possibilities, we used two different epigenetic modulators namely, 5-azacytidine (5-AZA) and suberoylanilide hydroxamic acid (SAHA) and determined their effect on cisplatin resistance. While treating with SAHA and cisplatin had no effect on H1993 cells (**Fig. 4D, right most panel**) or H23 (**Fig. 4D, left panel**), it had a significant additive effect on H2009 suggesting that these cells can become sensitive (**Fig. 4D, middle panel**). However, 5-AZA had no discernable effect (not shown) suggesting that epigenetic regulation of chromatin rather than specific cytosine residues in the DNA modulates cisplatin tolerance in the H2009 cells. Based on these criteria, H2009 qualify as cisplatin-tolerant (reversible) rather than resistant (irreversible) while H1993 may represent a truly resistant phenotype. Of note, the H1993 cells also carry amplification of MET, which may play a role in drug resistance. Taken together, these observations suggest that tolerance to cisplatin can be reversed unless the tolerant cells acquire a mutation making them irreversibly resistant (**Fig. 4E**).

### Mathematical modelling

To model such complex behavioral patterns, we tested various approaches using game theory and phenomenological modeling (Liao and Tlsty, 2014a; Liao and Tlsty, 2014b). Game theory approaches such as modified Lotka-Volterra and density games models (Novak et al, 2013) were developed, by introducing variable carrying capacity dependent on cellular composition (details in the **Supplementary text**). While such models were able to reproduce part of the observed phenomena, the agreement with the experimental trends was not overall satisfactory. Therefore, we developed a new theory based on dynamic phenotype switching in response to stress. The resulting model explained the growth behaviors for both monotypic and heterotypic experiments of various cellular compositions with and without cisplatin. As discussed below, the concept of phenotypic switching serves as a key survival strategy for cells within a multicellular system when faced with stressful conditions.

### Phenotype-switch model with stress response (PSMSR)

The detailed description of the PSMSR model is presented in the **Supplementary Text**. Briefly, we envision stress as a collection of factors in the cellular microenvironment that inhibit growth rate and increase cell death (schematic in **Fig. 5A**). The model presented here is based on the following assumptions: i) cellular growth leads to stress accumulation, ii) accumulated stress reduces growth, iii) tolerant cells are efficient in neutralizing stress, iv) stress triggers the switching of sensitive cells to the tolerant phenotype, and v) nutrients are assumed to be unlimited, since all the experiments were conducted in nutrient rich fresh media. Based on these assumptions, the growth rates of the sensitive and tolerant cells are given by equations 1 and 2, where S and T are the time varying populations of the sensitive and tolerant cells respectively.

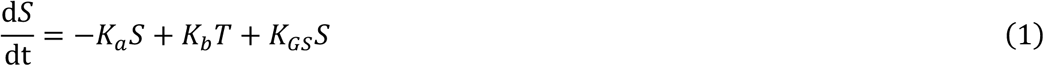

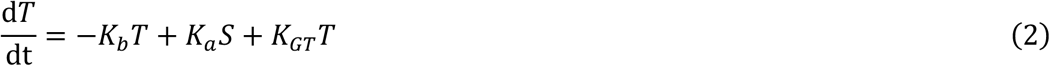

**Fig. 5:**
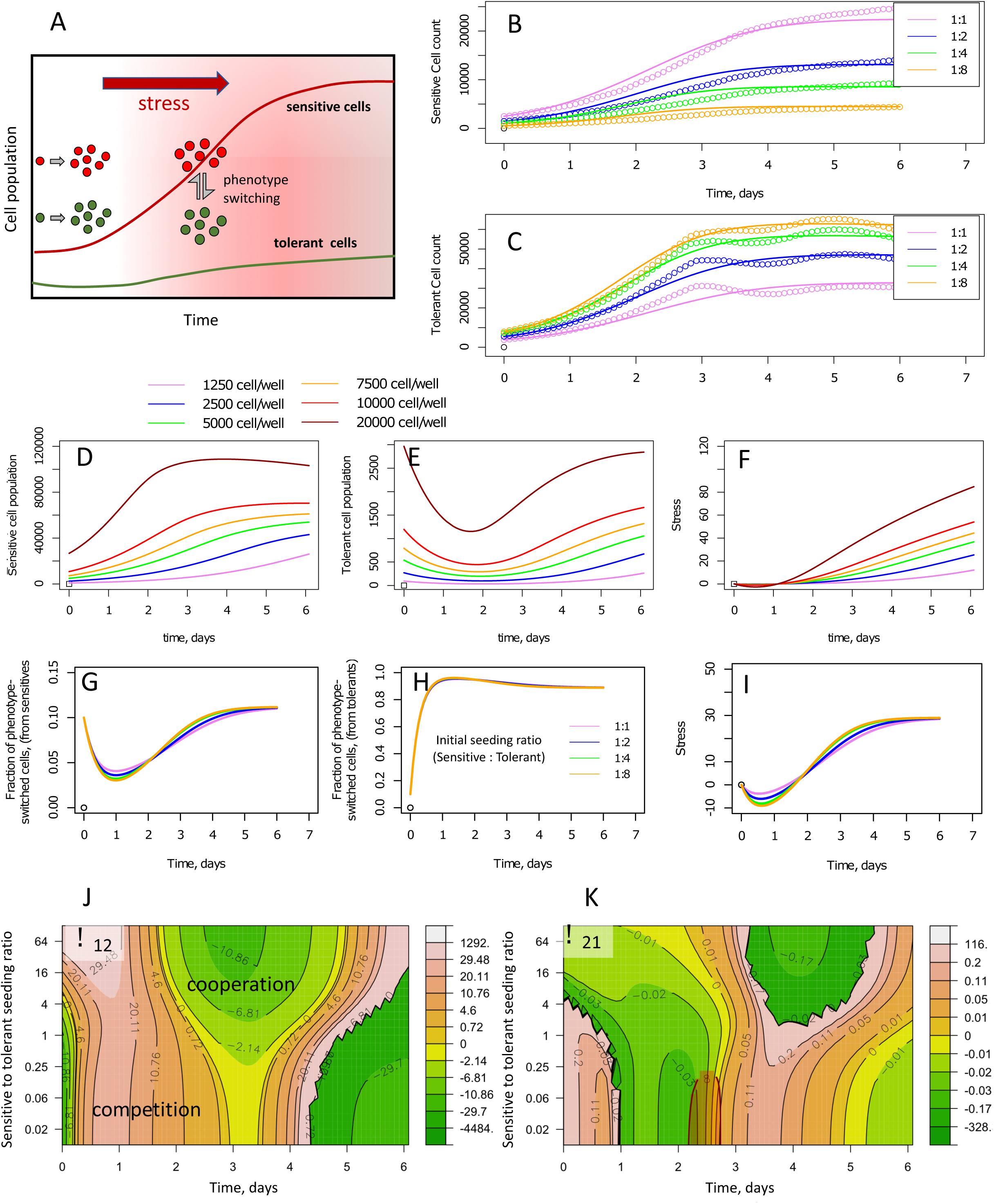
Cooperativity and stress response as described by the PSMSR model. (**A**) Schematic describing the PSMSR model; initially, the sensitive and the tolerant cells proliferate independently; as stress builds up, sensitive cells switch their phenotype to tolerant cells and vice versa; tolerant cells remove stress and maintain a small population, while enabling the sensitive cells to proliferate. (**B-C**) Fitting of the phenotype-switch model to the cellular growth curves of sensitive and tolerant cell populations, where the cells were mixed at different proportions and counting was started immediately; the colors represent the growth curves from different initial seeding proportions, as indicated in the legend (sensitive to tolerant cell seeding ratios); (**D-F**) predicted evolution of phenotype switching and stress in systems seeded with sensitive or tolerant cells alone; (**D-E**) populations of sensitive and (switched) tolerant phenotypes with time, when seeded with sensitive cells only; (**F**) stress as function of time; (**G-I**) predicted evolution of switched phenotypes and stress in mixed culture experiments, where cell growth was monitored immediately after mixing; (**G**) fraction of sensitive cells that have switched to the tolerant phenotype, as function of time; (**H**) fraction of tolerant cells that have switched to the sensitive phenotype, as function of time; (**I**) stress with time; colors are according to the initial seeding ratio of sensitive to tolerant cells as shown in the legend; the total cell population in each case was close to 5000; (**J-K**) evolving game strategy landscape of cellular population due to stress and phenotype switching; the heatmaps of time varying payoff values representative of inter-species competition/cooperation are shown as function of sensitive to tolerant seeding ratio; payoff values are derived by fitting the PSMSR model to the competitive Lotka-Volterra equations; orange areas in the maps represent competitive behavior, green areas represent cooperative behavior; (**J**) α_12_ representing the effect of tolerant cells towards the sensitive cells; (**K**) α_21_ representing the effect of sensitive cells towards the tolerant cells.

K_GS_ and K_GT_ are the stress-dependent effective growth rate constants (incorporating both proliferation and cell death) of the sensitive and tolerant cells respectively. K_a_ and K_b_ are the rate constants of switching from sensitive to tolerant phenotype and vice versa ^4^. We assume K_a_ to be stress-dependent and K_b_ to be constant (**Supplementary equations 13-15**).

Next we assume, that stress is predominantly generated by the fast growing sensitive cells at a rate proportional to the cell population, and neutralized by the tolerant cells. The resulting rate equation is given by:

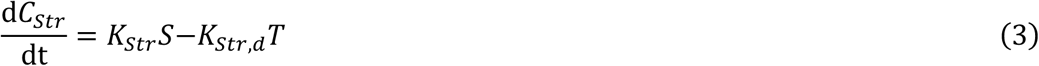

where *C*_*Str*_ stands for stress, and K_Str_ and K_Str,d_ are the rate constants of stress generation and removal respectively. Although, we have assumed simple linear relations between stress and growth or phenotype switching rates, in reality, these relations are likely to be more complex. Equations 1 and 2 were fitted to the experimentally measured cellular growth curves to obtain the parameters of the model (**Fig. 5B-C** and **Supplementary Fig. 8)** as described in the supplementary text.

### PSMSR and cisplatin response

To model the effect of cisplatin, we added a cisplatin dose-dependent cellular death rate term to equations 1 and 2 to obtain equations 4 and 5. Here AUC stands for “area under the curve”, which for a constant dose of cisplatin is simply the drug concentration S_drug_ x duration. Sigmoid (AUC) is a sigmoidal function (equation 6) that varies between 0 and 1, depending on the magnitude of AUC, which multiplied by the scale factor SCALE gives the cellular death rate. AUC_5_ and AUC_95_ represent the AUC values where 5% and 95% of the cisplatin death effect are achieved respectively. The SCALE, AUC_5_ and AUC_95_ parameters are specific for the sensitive and the tolerant cell types.

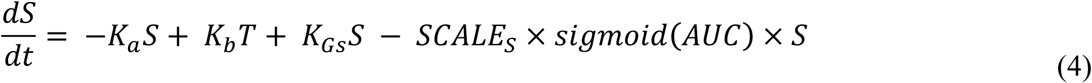

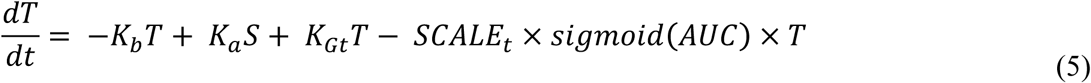

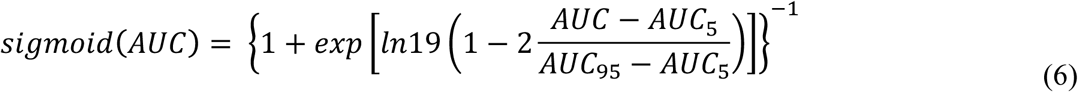

### PSMSR in pure and mixed cell populations

We first explored how the growth behaviors of the sensitive and the tolerant cells were affected by the coexistence of different phenotypes. In general, the phenotypic parameters for the heterotypic cultures were different in magnitude compared to these values in the monotypic cultures (**Supplementary Fig. 8J)**. This indicates an influence of the cellular phenotypes on each other. One striking result from the phenotype-switch model is that the growth rate constant for the tolerant phenotype in monotypic cultures is 7-10 times smaller than that of the sensitive phenotype (**Supplementary Fig. 8J)**. The slow growth of the tolerant cells is similar to the persister cells that impart drug resistance in bacterial population (Carvalho et al, 2019).

The PSMSR model has the ability to provide insights into the mechanism of phenotype switching in response to a changing microenvironment. Here, we have calculated the emergence of tolerant cell population from sensitive cells or vice versa in both monotypic (**Supplementary Fig. 9A, 9B, 9E** and **9F**) and heterotypic cultures **(Supplementary Fig. 10A, 10B, 10F** and **10G**). Of note, these cells are still experimentally detected as red or green, irrespective of their true phenotypes. Since stress is the driver for phenotype switching, the switched phenotype cells only appear once stress builds up in the system over time (**Supplementary Fig. 10**). In both pure and mixed cultures, the model predicts a rapid switch by the tolerant cells to the sensitive phenotype (up to 97%, **Supplementary Fig. 9E, 10B, 10D** and **10G**), while maintaining a low but steady tolerant population throughout the observations (**Fig. 5E, Supplementary Fig. 9B, 9F, Fig. 5A, 5F**). Thus, we have seen how the cancer cells use phenotype switching to maintain the overall fitness of the community under different stress levels. To maintain steady growth, stress must be mitigated, where switching to the tolerant phenotype pays off, since the tolerant cells are capable of neutralizing stress. However, the fraction of tolerant phenotype in the population is predicted to be low overall (10%, **Fig. 5G, Supplementary Fig. 10C** and **10H**). This may be due to a balance between the necessity for stress removal and the energy or other costs required to maintain the tolerant phenotype, which are not addressed in this work.

### Effects of phenotype switching and stress give rise to diverse game-theoretical strategies in mixed cell populations

Cancer cell behavior is widely studied using game theory-based models where the inter-species game strategies (competition and cooperation) are assumed to be constant throughout the growth regime. A familiar example of such a model is the competitive Lotka-Volterra equation (**Supplementary text, equations 1-2**), although more specialized models exist in the literature (Kareva and Karev, 2019). The phenotypic diversity available to cancer cells suggest that their game strategic landscape will be considerably complex, where the inter-species competition and cooperation are dynamically altered based on changing scenarios. We have explored whether the PSMSR model can capture this complex strategic landscape. By fitting the growth rates obtained from the phenotype switch model to the competitive Lotka-Volterra equations, we determined the inter-species competition parameters as function of time, for different sensitive to tolerant seeding ratios (**Fig. 5J and K**). We selected the Lotka-Volterra equation because it is the most familiar game theoretical model, although the approach can be applied to other models (**Supplementary Fig. 12**). Fig. 5J shows that the tolerant cells are initially competitive towards the sensitive cells, but they become cooperative after 2-3 days of growth. This coincides with the accumulation of stress (**Fig. 5G**) indicating that the microenvironment effects play a major role in altering the game strategies of the cancer phenotypes. Notably, the effect of the sensitive cells towards the tolerant cells (**Fig. 5K**) is significantly smaller in magnitude. In summary, the PSMSR model demonstrates the diverse strategic landscape explored by the sensitive and tolerant cells under varying cell population and stress levels.

### PSMSR model demonstrates the effectiveness of the intermittent cisplatin therapy

The supplementary text describes the details of the cisplatin effect modeling. The best agreement between the growth trends predicted by the PSMSR model and the experiments (**Supplementary Fig. 7A, C, Fig. 6B-C and Supplementary Fig. 7B, D**) was obtained when community cooperation such as phenotype switching and stress removal were turned off. This implies that high cisplatin levels trigger the tolerant cells to focus on their own survival, similar to other social communities where imminent danger promotes self-survival. Also at high cisplatin levels, the phenotype switching to sensitives is detrimental to the survival of the community. Together, these observations are indicative of the adaptability of the cancer cells for survival in adverse environments. In Fig. 6D, the magnitudes of the SCALE parameter quantify the difference in cisplatin sensitivity between the sensitive and the tolerant phenotypes. Curiously, the sensitive cells appear to be more cisplatin-tolerant after co-culture with the tolerant cells, as shown by the lower SCALE parameter after three weeks of co-culture. This is in line with our observations from the spatiotemporal behavior of the sensitive cells that indicate learning from the tolerant cells.

**Fig. 6:**
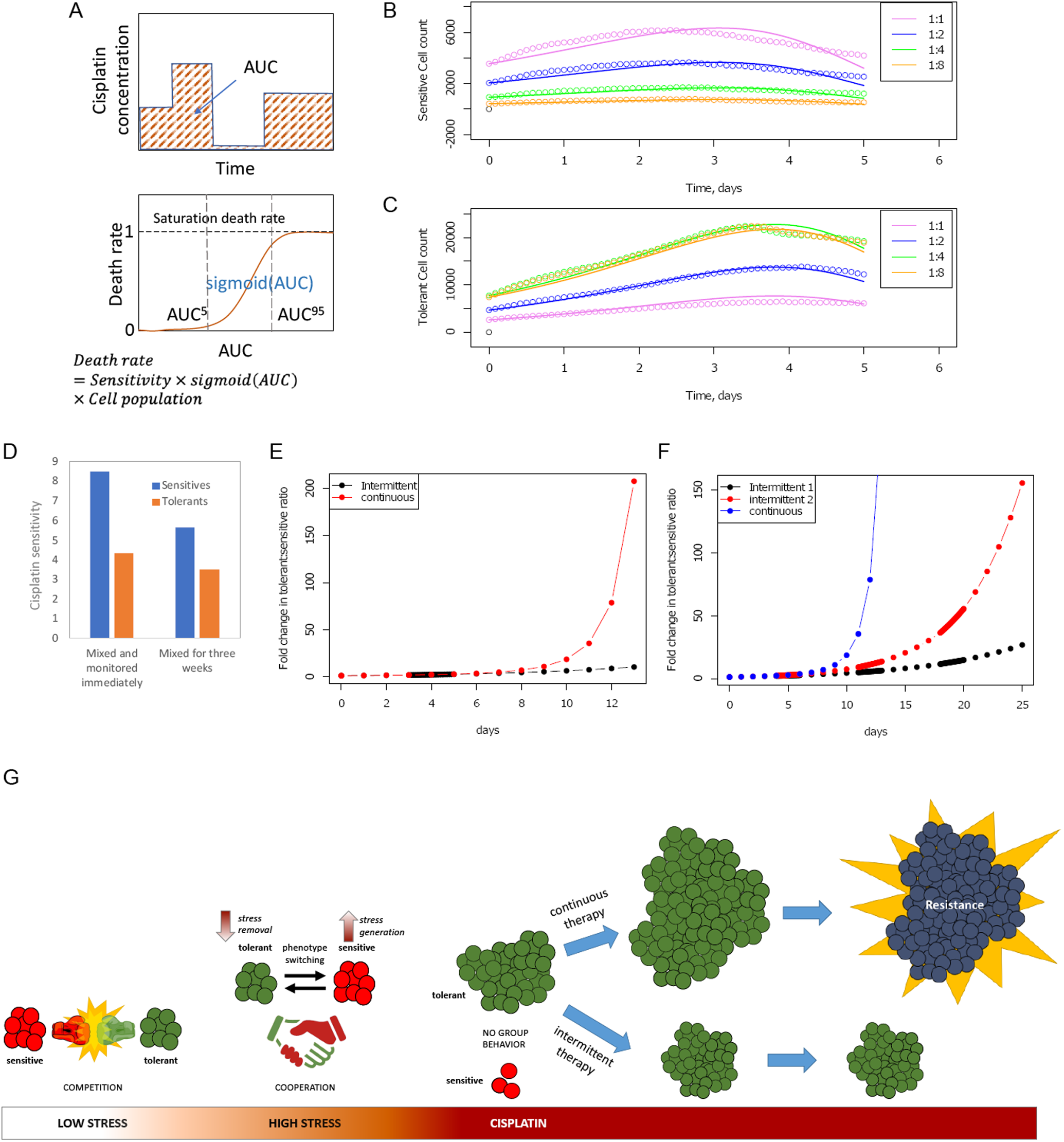
Mathematical model for cisplatin resistance. (**A**) Schematic demonstration of AUC and cellular death rate as function of AUC; (**B-C**) fitting of the experimental growth data where the cells were co-cultured for three weeks; B: sensitive cells; C: tolerant cells; circles and lines represent the experimental and fitted trends respectively; (**D**) SCALE parameter as measure of cisplatin sensitivity for the sensitive and the tolerant cells; (**E-F**) simulation of intermittent and continuous cisplatin treatment according to the protocols described in Fig. 3; the initial sensitive to tolerant cell ratio was set to 4:1 with a total cell population of 50,000. (**G**) An illustrative model depicting the presence (and absence) of group behavior among sensitive and tolerant cells under varying conditions of stress and effects of continuous versus intermittent therapy.

Using the PSMSR model, we simulated the cisplatin dose cycles as described in **Fig. 3**. Similar to the experimental observations, the tolerant cell proportion increased with time in the continuous therapy, while it remained relatively small and increased at a slower rate during the intermittent cycles (**Fig. 6E-F**). Also, the intermittent 2 cycles of cisplatin dosage created more tolerant cell population than the single cycle, in agreement with the experiments **Fig. 4**. While the actual magnitudes of the tolerant cell population in the simulations are different than in the experiments, the qualitative behaviors are in agreement. The quantitative difference between the predictions and the experiments could be attributed to the growth attenuation due to confluency that is accounted for by the model. Overall, these simulations show that the PSMSR model is able to qualitatively reproduce the drug induced behavior of the cancer cell population.

## Discussion

Several studies have applied evolutionary game theory to cancer (Wu et al, 2014; Wu et al, 2015; Zhang et al, 2017; You et al, 2017; Swierniak et al, 2018; Kareva and Karev, 2019; Kaznatcheev et al, 2019; Stanková et al, 2019; West et al, 2019) but as far as we are aware, the adaptive strategies that drug naïve and drug tolerant or resistant NSCLC cancer cells adopt in response to environmental perturbations have not been investigated. A recent paper by Kaznatcheev et al (2019) is the only other study where the authors developed an assay to measure effective evolutionary games in co-cultures of isogenic NSCLC cells that are sensitive and resistant to the anaplastic lymphoma kinase inhibitor alectinib. In contrast, in the present study we investigated a broad spectrum chemotherapy (cisplatin) and naturally occurring cisplatin-sensitive and tolerant NSCLC cell lines and developed the PSMSR model to model the experimental observations. Our rationale for not choosing isogenic cell lines is predicated on new thinking in the field. It is generally held that genomic instability underlies many cancers and generates genetic variation that drives cancer initiation, progression, and therapy resistance. However, in contrast to the prevailing wisdom that mutations occur purely stochastically at constant, gradual rates, emerging evidence strongly indicates that many organisms and human cancer cells in particular, possess mechanisms of mutagenesis that are upregulated by stress responses. Thus, such transient, genetic diversity bursts can propel evolution, especially when cells are poorly adapted to their environments (when stressed) (Fitzgerald et al, 2017) underscoring the limitations of the isogenic strains. It is quite plausible that such (artificially) stress-induced mutations could also potentially confound cell behaviour.

Our results lend further support to previous suggestions (Zhang et al, 2017; Swierniak et al, 2018; Cunningham et al, 2018; Stanková et al, 2019) that, knowledge regarding the ecological and evolutionary strategies exploited by cancer cells, could be used to delay or even prevent drug tolerance and resistance (**Fig. 4**). We demonstrated that the growth behaviour of the sensitive and tolerant cells can be explained by invoking dynamic phenotypic switching in the presence of stress. Consistent with this assumption, our previous studies (Mooney et al, 2016; Mohanty et al, 2019) and several other studies (Gupta et al, 2011; Goldman et al, 2015) revealed that the two cell types can stochastically switch their phenotypes via a non-genetic mechanism. Our experimental observations and the insights gained from applying mathematical modelling illustrate the complex behavioral landscape of the cancer cells, where the payoff strategies are dynamically evolving via phenotypic plasticity and environmental pressure. Such complex inner level traits are reflected in macro properties such as the carrying capacity, which we find to be cell frequency-dependent.

The PSMSR model demonstrated that stress, which is a consequence of cellular growth, promotes phenotype switching to create more tolerant cells. These cells then partly neutralize the stress to allow the fast-growing sensitive cells to proliferate, thereby sustaining the carrying capacity of the system. Thus, the resulting carrying capacity is a function of both stress and the level of tolerant cells in the system. Interestingly, the behavior of the tolerant cells appeared to be beneficial to the overall community, not just themselves, as they helped the sensitive cells to grow by removing stress and adopting a slow proliferation rate, so as to not compete for limited resources. It should be noted, that both sensitive and tolerant cells have similar doubling times as is reflected in their intrinsic growth rate constants. However, due to the high phenotype switch rate (parameter K_b_, **Supplementary Fig. 8J**), a large portion of emerging tolerant cells may switch to the sensitive phenotype, either by keeping the tolerant phenotype population low. Thus, the tolerant cells appeared to exhibit an ‘altruistic’ behaviour, which is the result of their dynamic phenotype switching behavior. In contrast, the sensitive cells appeared less likely to change their phenotype, even after exposure to the tolerant cells for three weeks, as is evident in their low equilibrium constant of switching to tolerant cells (parameter K_0_, **Supplementary Fig. 8J**). Taken together, the behavior of the tolerant cells provides novel insights into the phenotypic traits in cancer that emerge due to survival pressure as well as the cost-benefit basis of such evolution. Of note, when co-cultured for three weeks prior to starting the experiment, both the sensitive and tolerant cell types had equal opportunity to switch their strategy; either they could have reduced or increased their proliferation rate to compete with each other, but they followed an unexpected path where the sensitive cells remained unaffected or the tolerant cells reduced their proliferation rate. This strategy could be helpful because most of the genotoxic drugs used for chemotherapy target actively dividing cells. This behavior of tolerant cells resembles phenotypic switching behavior reminiscent of persisters that are well known in microbial systems (Brauner et al, 2016) and more recently being recognized in cancer (Dawson et al, 2011; Hata et al, 2016; Jolly et al, 2018; Vallette et al, 2019).

Consistent with this prediction, several studies (Kumar et al, 2019; Jolly and Celià-Terrassa, 2019) including our own studies (current and Mohanty et al 2019) reveal that, tumor cells can switch phenotypes in response to the presence/absence (concentrations) of the drug. Therefore, contrary to the dogma that drug resistance is due to the selection of a pre-existing mutant clone (Bozic and Nowak, 2014), our study supports the deluge of emerging evidence indicating that non-genetic mechanisms may also help cancer cells evade drug treatment (Ahmed and Hass, 2028; Salgia and Kulkarni, 2018; Bell and Gilan, 2019; Xu et al, 2020; Hartman et al, 2020; Gunnarsson et al, 2020). It now appears that cancer cells employ non-genetic, genetic and a combination of genetic and non-genetic mechanisms of drug resistance (Chisholm et al, 2015) underscoring the cancer cell’s remarkable resilience. Furthermore, it is also suggested that there may be an interaction between genetic and non-genetic adaptation that defines ‘the path of most resistance’, which outlines the variables that dictate whether cancers adapt through genetic and/or non-genetic resistance (Bell and Gilan, 2019).

The present study suggests that intermittent rather than continuous chemotherapy may result in better outcomes in lung cancer. As far as we are aware, such an approach, although investigated in some other solid tumors (Cunningham et al, 2015; Zhang et al, 2017), has never been looked into in lung cancer although, the observations by Kaznatcheev et al (2019) are inferred to support the concept that treatment influences the character of the resistance response in cancer cells (Staňková K, 2019). Such a treatment approach may not cure the patient of the disease, but it would at least result in stable disease that can be managed while sparing the patient of undesirable effects of excessive chemotherapy. On the contrary, current chemotherapy protocols may encourage cancer cells to ‘anticipate’ (Saigusa et al, 2008) sustained chemotherapy. As a result, it is possible that they permanently switch to a resister phenotype via a genetic mutation (Fitzgerald et al, 2017; Gunnarsson et al, 2020) and therefore, worsen disease prognosis by promoting the selection of irreversible and truly resistant cells that emerge from a potentially reversible, ‘tolerant’ population (**Fig. 4**). Thus, preventing phenotype switching using inhibitors can suppress the emergence of temporary drug tolerance and enhance the efficiency of intermittent therapy. Designing such interventions that augment conventional therapy requires insights into the dynamic group behavior of cancer cells in mixed cell environments, as has been done in the current study. Additional studies that compare truly ‘resistant’ (H1993) cells to sensitive/tolerant cells, and studies employing 3D cultures including components of the tumor microenvironment such as cancer-associated fibroblasts, that are currently underway in our laboratories, should provide further insight on how group behaviour impacts drug resistance in cancer.

## Materials and Methods

### Cell lines and reagents

Cell lines (H23, H2009, H1993) were obtained from American Type Culture Collection (ATCC) (Manassas, VA, USA) and cultured in RPMI 1640 medium (Corning) supplemented with 10% fetal bovine serum (FBS), L-glutamine (2 mM), penicillin/streptomycin (50 U/ml), sodium pyruvate (1 mM), and sodium bicarbonate (0.075%) at 37°C, 5% CO_2_. Cisplatin was provided by City of Hope National Medical Center clinics (Duarte, CA, USA). Puromycin was purchased from Thermo Fisher Scientific (Waltham, MA, USA). Suberoylanilide hydroxamic acid (SAHA) was purchased from Selleck Chemicals (Houston, TX, USA).

### Live cell imaging and analysis

Cell lines H23 and H1993 were stably transfected with NucLight Red Lentivirus (Essen Bioscience, Ann Arbor, MI, USA) to express nuclear mKate2, a red fluorescent protein (RFP), and H2009 was stably transfected with NucLight Green Lentivirus to express nuclear green fluorescent protein (GFP). Stable clones were selected with puromycin (1 μg/ml). Live cell images were acquired in real time using the IncuCyte Live Cell Imaging System (Essen Bioscience). Cell counting masks were generated using the IncuCyte software to perform data analysis (Mohanty et al, 2019).

### Monotypic/heterotypic culture cell proliferation and drug sensitivity assay

For 1:1 heterotypic cultures, 2.5×10^5^ H23 (cisplatin-sensitive) cells and 2.5×10^5^ H2009 (cisplatin-tolerant) cells (5×10^5^ total cells) were plated on a 10 cm dish. For heterotypic cultures with different sensitive : tolerant seeding ratios (1:2, 1:4, 1:8, 2:1, 4:1, 8:1), numbers of each cell type were adjusted accordingly and plated with 5×10^5^ total cells per plate. Cultures were maintained in 37°C, 5% CO_2_ for 3 weeks and passaged 1:5 when confluent (every 4-5 days). After 3 weeks, 5×10^3^ cells from each culture were seeded on a 96-well plate and allowed to adhere overnight. Cisplatin was added at the indicated concentration. Cell proliferation was monitored in real time using the IncuCyte Live Cell Imaging System (Essen Bioscience), and data was collected every 2 hours. Data analysis was performed using the IncuCyte software using a red and green fluorescence mask to accurately count each cell type. Statistical significance was measured using ANOVA test (two or one-way) and t-test.

### Conditioned medium assay

Complete growth medium was added to 4×10^6^ cells on a 10 cm dish to condition medium. After 24 hours, conditioned medium was collected in a centrifuge tube and spun down at 2500 RPM for 10 mins. Afterwards, conditioned medium was added to 5×10^3^ cells seeded on a 96-well plate. Cell proliferation was monitored in real time using the IncuCyte Live Cell Imaging System (Essen Bioscience), and data was collected every 2 hours. Data analysis was performed using the IncuCyte software using a red and green fluorescence mask to accurately count each cell type.

### Cell clustering and nearest neighbor analysis

Sensitive cells and tolerant cells were seeded in monotypic or 1:1 heterotypic (5×10^3^ total cells) cultures on a 96-well ImageLock plate (Essen Bioscience). Growth conditions were: complete medium (10% serum), low serum medium (0.5% serum), or cisplatin (1 μM). Cell clustering was monitored in real time using the IncuCyte Live Cell Imaging System (Essen Bioscience), and data was collected every 20 minutes. Data analysis was performed using the IncuCyte software using a fluorescence mask to accurately count each cell type. QuPath 0.2.0M8 software (https://github.com/qupath/qupath/wiki/Citing-QuPath) was used to first segment cells using the Cell Detection function, then connectivity between cell centroids within a distance of 35 microns was determined by calculating Delaunay Cluster Features (Bankhead 2017). Cells in clusters gain additional measurements based on their neighbors within the cluster. The cells highlighted in white in the image (Supplementary Fig 5A) have 7 neighbors (number of connected cells not including itself) as seen on the right image. Similarly, the cells that do not have any neighbors will be color-coded as black and have number of neighbors as 0; cells with increasing numbers of neighbors are color-coded according to the scale shown, starting with blue cells having one neighbor. To find the nearest neighbor distance, the minimum distance was changed from 35 microns to the size of the entire image, generating one large cluster. Each cell in that large cluster then gained a measurement representing the distance to the nearest cell and the average value of that could be reported for a given image.

### Cell migration assay and tracking analysis

Sensitive cells were seeded in monotypic or 1:1 heterotypic culture with tolerant cells (5×10^3^ total cells) on a 96-well ImageLock plate (Essen Bioscience). Growth conditions were: complete medium (10% serum), low serum medium (0.5% serum), or cisplatin (1 μM). Cell migration was monitored in real time using the IncuCyte Live Cell Imaging System (Essen Bioscience), and data was collected every 20 minutes. Data analysis was performed using the IncuCyte software using a fluorescence mask to accurately count each cell type. Grayscale TIFF images were exported from IncuCyte and imported as a series into ImagePro Premier. Calibration values were added so that speed and distance traveled could be calculated using microns. Each sequence of images was further subdivided into three bins based on cell mean intensity, and tracks were generated by following those cells using the Tracking function. The binning reduced the number of times cells would run near or through each other and exchange tracks. The low, medium, and high intensity track data were then merged per series to produce summary values.

### RNA sequencing

To identify the effect of the sensitive or tolerant cell on gene expression, sensitive cells as a monotypic culture or a 1:1 heterotypic culture with tolerant cells were cultured for a period of 5 days. After 5 days, with assistance from the Analytical Cytometry Core at City of Hope using the BD FACSAria II cell sorter (BD Biosciences, San Jose, CA, USA), the mixed population was separated based on the fluorescence tag, either GFP or RFP. Total RNA was extracted from these cells with RNeasy Mini Kit (Qiagen, Hilden, Germany) and sent to the Integrative Genomics Core at City of Hope for exome sequencing, with 40 million reads per sample. The sequencing information was used for comparative analysis between the monotypic sensitive versus monotypic tolerant and heterotypic sensitive versus monotypic tolerant. Ingenuity Pathway Analysis (IPA) was used to identify the pathways that showed significant change. The IPA platform was used to determine the changes in expression of genes involved in cell movement.

### SAHA treatment and drug sensitivity assay

Cells were (3×10^5^) plated on a 6 cm dish and allowed to adhere overnight. Fresh medium containing SAHA (0.25 μM/0.5 μM) was added every 24 hours for 3 days. After 3 days, 5×10^3^ cells were seeded on a 96-well plate and allowed to adhere overnight. Cisplatin (5 μM) was added, and cell proliferation was monitored in real time using the IncuCyte Live Cell Imaging System (Essen Bioscience). Data was collected every 2 hours, and analysis was performed using the IncuCyte software using a red and green fluorescence mask to accurately count each cell type.

## Acknowledgments

MKJ was supported by Ramanujan Fellowship (SB/S2/RJN-049/2018) awarded by Science and Engineering Research Board, Department of Science and Technology, Government of India. GR was supported by JC Bose National Fellowship, Government of India and UGC Centre of Advanced Study.

## Author Contributions

A Nam performed experiments, analyzed data, and wrote the manuscript, AM conceptualized the idea, performed experiments, analyzed data, and wrote the manuscript, SB conducted the mathematical analysis and wrote the manuscript, SK conducted the mathematical analysis and wrote the manuscript, SA formulated the analytical approach, analyzed data, and reviewed the manuscript, KH analyzed data, A Nathan analyzed data, GR formulated the analytical approach and reviewed the manuscript, EM reviewed the manuscript, HL reviewed the analytical approach and reviewed the manuscript, MKJ formulated the analytical approach and reviewed the manuscript, PK conceptualized the idea and wrote the manuscript, RS conceptualized the idea and reviewed the manuscript

## Supplementary Figure Legends

**Supplementary Fig. 1.** *Generating heterotypic cultures in 1:1 ratio and increasing T:S ratios (2:1, 4:1, 8:1).* (**A**) Cisplatin-sensitive cells (H23) and cisplatin-tolerant cells (H2009) stably express nuclear red fluorescent protein (RFP) and green fluorescent protein (GFP), respectively. (**B**) Schematic representation of the experimental design of co-culturing sensitive (S) and tolerant (T) cells in T:S ratios of 1:1, 2:1, 4:1, and 8:1 for 3 weeks and collection of data points.

**Supplementary Fig. 2.** *Proliferative behavior of sensitive (S) and tolerant (T) cells mixed in increasing T:S seeding ratios 12 hours prior to the start of the experiment.* (**A**) Schematic representation of the experimental design of mixing cells in various T:S ratios (1:1, 2:1, 4:1, and 8:1) 12 hours prior to cisplatin treatment and real-time monitoring of cell proliferation. (**B**) Proliferation rate of sensitive cells (red) and tolerant cells (green) in heterotypic culture over the course of 144 hours. (**C**) Proliferation rate of sensitive cells (red) and tolerant cells (green) in heterotypic culture treated with 5 μM cisplatin over the course of 144 hours. (**D**) Change in tolerant/sensitive cells ratio with (orange) and without (black) 5 μM cisplatin over the course of 144 hours. Statistical significance information can be found in **Supplementary Table 5**.

**Supplementary Fig. 3.** *Proliferative behavior of sensitive (S) and tolerant (T) cells grown in heterotypic cultures of increasing S:T seeding ratios for 3 weeks.* (**A**) Schematic representation of the experimental design of co-culturing sensitive (S) and tolerant (T) cells in S:T ratios of 1:1, 2:1, 4:1, and 8:1 for 3 weeks and collection of data points. (**B**) Proliferation rate of sensitive cells (red) and tolerant cells (green) in heterotypic culture over the course of 144 hours. (**C**) Proliferation rate of sensitive cells (red) and tolerant cells (green) in heterotypic culture treated with 5 μM cisplatin over the course of 144 hours. (**D**) Change in tolerant/sensitive cells ratio with (orange) and without (black) 5 μM cisplatin over the course of 144 hours. Statistical significance information can be found in **Supplementary Table 6**.

**Supplementary Fig. 4.** *Clustering analysis of sensitive and tolerant cells grown in monotypic and heterotypic cultures under various conditions.* (**A**) Images showing the clustering of monotypic sensitive and tolerant cells after 24 h of culturing in 10% serum. Each cell was color coded to reciprocate its number of nearest neighbors. Cells colored white have > 7 number of nearest neighbors, and the cells colored black have 0 nearest neighbors. (**B**) The distribution frequency of cells with nearest neighbors ranging from 0-8. The solid green bar graph on the left represents nearest neighbors for tolerant cells in monotypic culture, and the light green graph represents distribution in heterotypic culture. Under heterotypic culture the distribution frequency shifts towards left, with less number of nearest neighbors. (**C**) Similarly, the nearest neighbors were also calculated for the cells in presence of cisplatin for 24 h. The frequency of cells with fewer neighbors increases more in heterotypic cultures.

**Supplementary Fig. 5.** *Migratory behavior of sensitive cells affected in heterotypic culture as compared to monotypic culture.* (**A**) Graphical representation of migration tracks of 200 sensitive cells in monotypic or heterotypic cultures (1:1 sensitive versus tolerant) under 10% serum (normal), 0.5% serum, or cisplatin. (**B**) Schematic representation of the gene expression analyzed between sensitive (monotypic) versus tolerant (monotypic), in the blue circle and sensitive (heterotypic) versus tolerant (monotypic) in the orange circle. The overall outcome of 132 genes in the monotypic culture led to decrease in cell movement and on the same place outcome of 973 genes in heterotypic culture caused higher cell movement. (**C**) There are set of 9 genes which were common in both comparisons. The heat map representing the log2 fold change in their gene expression from monotypic to heterotypic culture.

**Supplementary Fig 6.** *Fits of simulation (dashed lines) to the mean population size data (solid lines) for different conditions and models are shown.* (**A-C**) for 3-week co-culture data with interaction models representing Predation by Sensitive cells, Predation by Resistant cells and Symbiosis respectively.

**Supplementary Fig. 7**. *Fits of simulation (dashed lines) to the mean population size data (solid lines) for different conditions.* (**A-B**) Individual culture data for sensitive and tolerant cells, respectively. (**C-D**) 3-week co-culture data with interaction models representing Competition for sensitive and tolerant cells, respectively. (**E** and **H**) Model fit for sensitive cell population in mixed culture without and with cisplatin, respectively. (**F** and **I**) Model fit for tolerant cell population in mixed culture without and with cisplatin, respectively. (**G** and **J**) shows the fitting error in mixed culture without and with cisplatin, respectively. (**K** and **L**) show payoff values for mixed culture without and with cisplatin, respectively.

**Supplementary Fig. 8**. *Fitting of the PSMSR model to the cellular growth curves*; (**A**) sensitive cells only; (**B**) tolerant cells only; the curves are colored according to the initial seeding population; circles represent the cell populations at different time points as means of three replicates; lines represent the fitted trends; (**C**) estimated fitting error for sensitive only (blue) and tolerant only (orange) populations; (**D, E**) sensitive and tolerant cell populations, where the cells were mixed at different proportions and counting was started immediately; the colors represent the growth curves from different initial seeding proportions, as indicated in the legend (sensitive to tolerant cell seeding ratios); (**F**) fitting error for sensitive and tolerant cell growth in the mixed culture; (**G-I**) equivalent results for the mixed cell populations, where the cells were cultured together for three weeks before monitoring growth; (**J**) parameter values for the various cell systems; the parameters that significantly differed for the 3 weeks cultured population compared to the other systems are highlighted in green (increase) or red (decrease).

**Supplementary Fig. 9**. *Predicted evolution of phenotype switching and stress in systems seeded with sensitive or tolerant cells alone*; (**A-B**) populations of sensitive and (switched) tolerant phenotypes with time, when seeded with sensitive cells only; (**C-D**) ratio of sensitive to tolerant phenotype (C) and stress (D) as functions of time; (**E-F**) populations of (switched) sensitive and tolerant phenotypes when seeded with tolerant cells only; (**G-H**) ratio of sensitive to tolerant phenotype (G) and stress (H) as functions of time; colors represent different seeding populations.

**Supplementary Fig. 10**. *Predicted evolution of switched phenotypes and stress in mixed culture experiments*; A-E represent systems where growth was monitored immediately after mixing; F-J represent systems where growth was monitored three weeks after mixing; (**A, F**) tolerant cell population with time that have switched their phenotype from sensitive cells; (**B, G**) sensitive cell population with time that have switched phenotype from tolerant cells; (**C, H**) fraction of sensitive cells that have switched to the tolerant phenotype, as function of time; (**D, I**) fraction of tolerant cells that have switched to the sensitive phenotype, as function of time; (**E, J**) stress with time; colors are according to the initial seeding ratio of sensitive to tolerant cells as shown in the legend; the total cell population in each case was close to 5000.

**Supplementary Fig. 11.** *Integration between the PSMSR and competitive Lotka-Volterra models*; the Lotka-Volterra model parameters are derived by fitting the LV equation to the growth rate of sensitive and tolerant cells as calculated by the PSMSR model over different time windows, initial seeding of 7500 total cells with different sensitive to tolerant mixing ratios; the graphs show the comparison between growth rates calculated from both models; rates from PSMSR model are in black; rates from LV equation fitting are in red; the best fitted parameter values and fitting error are shown in the legend of each plot.

**Supplementary Fig. 12.** *Integration between the PSMSR and Density-games based models*; the density-games model parameters are derived by fitting the mLV equation (shown in D) to the growth rate of sensitive and tolerant cells as calculated by the PSMSR model over different time windows, initial seeding of 5000 total cells with a sensitive to tolerant mixing ratio of 1:1; (**A-C**) the agreement between the growth rates from the best fitted mLV model to the PSMSR model for sensitive and tolerant cells over time windows of 12 hours, 4 and 6 days, respectively; the best fitted parameter values and fitting error are shown in the legend of each plot; (**D**) the mLV equations from the Density-games model that were fitted to the PSMSR model; (**E**) comparison between the mLV parameters obtained from fitting to the experimental data to the ones derived by fitting to the PSMSR model for an integration time window of 4 days.

**Supplementary Table 1.** *Ratio comparison of tolerant to sensitive cells at time of seeding and after 12 hours or 3 weeks of co-culture.* Rows 1-4 show the increase in tolerant population after a period of 12 h, and rows 5-8 show the changes after three weeks. Rows 9-12 show the changes in ratio when sensitive cells were seeded at higher number compared to tolerant cells. The comparison of rows 4-8 to rows 10-12 indicate that if a heterogeneous population has more sensitive cells initially, they will grow faster and dominate over the tolerant cells.

**Supplementary Table 2.** *Statistical significance for Fig. 1B-D*. Two-way ANOVA test was performed to determine statistical significance of last time point (144 h).

**Supplementary Table 3.** *Statistical significance for Fig. 2A*. Two-way ANOVA test was performed to determine statistical significance of last time point (72 h).

**Supplementary Table 4.** *Statistical significance for Fig. 4A-C*. Wilcoxon two-tailed test was performed to determine statistical significance.

**Supplementary Table 5.** *Statistical significance for Supplementary Fig. 2B-D.* Two-way ANOVA test was performed to determine statistical significance of last time point (144 h).

**Supplementary Table 6.** *Statistical significance for Supplementary Fig. 3B-D.* Two-way ANOVA test was performed to determine statistical significance of last time point (144 h).

