## Supplemental Text for "Suppressing chemoresistance in lung cancer via dynamic phenotypic switching and intermittent therapy"

**Supplementary text**

In this work, we have experimentally studied the growth of chemo-sensitive and tolerant cancer cells in mixed culture experiments with or without the presence of cisplatin. We have modeled the cellular growth using different mathematical models to extract the inherent principles of cellular dynamics. Using game theory based on modified Lotka-Volterra and density like game models, we have analyzed the competition and cooperation between the sensitive and the tolerant cell species when they are mixed together. We have also studied the cellular growth in pure cell cultures, where only a single 'player' is present where the game theory models are less suitable. To address this, we have therefore used a model based on rate-kinetics and phenotype switching that explains the role of environmental stress on the phenotypic evolution. The rate-kinetics based model demonstrates how the finer level effects such as stress and phenotypic plasticity shape the broader strategies employed by the cells, such as competition and cooperation.

Mathematical modeling based on game theory to analyze population dynamics measurements is often employed to elucidate complex dynamic processes in biological systems, including cancer (Wu et al, 2014; Wu et al, 2015; You et al, 2017; Zhang et al, 2017; Swierniak et al, 2018; Kareva and Karev, 2019; Kaznatcheev et al, 2019; Stanková et al, 2019). Game theory, particularly evolutionary game theory (EGT) can provide insights into the cooperation/competition among different cells in the tumour microenvironment, and into the design of potential therapies that disrupt these interactions (Archetti and Pienta, 2019). EGT has been used as a tool (Liao and Tlsty, 2014a; Liao and Tlsty, 2014b) to mathematically model various facets of cancer cells such as apoptosis, angiogenesis, tumor invasion, and treatment response in different cancer types (Gatenby and Maini, 2003). Two player games with pure or mixed strategies (Spaniel, W, 2014) as in the case of Hawk-Dove (HD) or pure strategies with Prisoner's Dilemma (PD) have been employed to study tumor cell evolution in the past (Hurlburt et al, 2018) to account for a more realistic population-based investigation. For complex cellular processes to constantly adapt to changing environments individuals would need to adopt competitive, submissive or mixed strategies as in the case of the HD game for some period, and adopt pure strategies such as co-operative and anti-cooperative strategies as in PD at other times (Kareva and Karev, 2019).

Our results lend further support to previous suggestions (Zhang et al, 2017; Swierniak et al, 2018; Cunningham et al, 2018; Stanková et al, 2019) that, knowledge regarding the ecological and evolutionary strategies exploited by cancer cells, could be used to delay or even prevent drug tolerance and resistance (**Fig. 4**). We demonstrated that the growth behaviour of the sensitive and tolerant cells can be explained by invoking dynamic phenotypic switching in the presence of stress. Consistent with this assumption, our previous studies (Mohanty et al, 2019) and several other studies (Mooney et al, 2016; Gupta et al, 2011; Goldman et al, 2015) revealed that the two cell types can stochastically switch their phenotypes via a non-genetic mechanism. Our experimental observations and the insights gained from applying mathematical modelling illustrate the complex behavioral landscape of the cancer cells, where the payoff strategies are dynamically evolving via phenotypic plasticity and environmental pressure. Such complex inner level traits are reflected in macro properties such as the carrying capacity which we find to be cell frequency-dependent. Traditional game theory models such as the unmodified LV, are ill equipped to handle such complexities. We therefore came up with a novel empirical growth equation based on density games model that qualitatively explained the observed growth behavior. However, unlike the traditional LV model which can be easily understood in terms of inter-species competition and fixed resources, the empirical assumptions in our game theoretical model such as the time varying carrying capacity are not immediately interpretable. Therefore, we modeled the growth behavior using an orthogonal approach where we assumed that a part of the genetically tagged sensitive and tolerant cells switch to their counter phenotypes in response to stress. This leads to the phenotype-switch model which not only explained the growth behavior of the cells but also demonstrated how a fixed set of rate kinetic parameters (one each for the sensitive and tolerant phenotypes) led to the time and seeding-dependent game-theoretical properties obtained from the density game-like model.

The phenotype-switch model demonstrated that stress which is a consequence of cellular growth, promotes phenotype switching to create more tolerant cells. These cells then partly neutralize the stress to allow the fast growing sensitive cells to proliferate, thereby sustaining the carrying capacity of the system. Thus, the resulting carrying capacity is a function of both stress and the level of tolerant cells in the system. This explains why the carrying capacity could be time and seeding composition-dependent since stress varies with time and the level of tolerant cells in the system, that appear through phenotypic switching, is dependent on the initial seeding. The phenotype switch model, therefore, helps to explain the basis of the most novel features of the game-theoretic models, emphasizing the importance of such a multi-level approach to the current problem.

We used EGT to model the temporal interactions of a population of cisplatin-sensitive and cisplatin-tolerant cell lines grown in 2D cultures. The Lotka-Volterra equation, particularly the competitive Lotka-Volterra equation (see Supplement section for a description), is the equation of choice to begin with. The form is similar to the Lotka-Volterra equations for predation in that the equation for each species has one term for self-interaction and one term for the interaction with other species. In the equations for predation, the base population model is exponential. For the competition equations, the logistic equation is the basis.

In the case of 2D, the competitive Lotka-Volterra (LV) equation with 2 players that have payoffs defined by a competitive matrix A=($\alpha_{ij}$) (Zhang et al, 2017) is typically defined as follows:

$$\frac{dS}{dt} = r_1\, S\left(1 - \left(\frac{\alpha_{11}S + \alpha_{12}T}{K_1}\right)\right) \quad (1)$$

$$\frac{dT}{dt} = r_2\, T\left(1 - \left(\frac{\alpha_{21}S + \alpha_{22}T}{K_2}\right)\right) \quad (2)$$

where $S$ refers to sensitive cell density and T refers to tolerant cell density, $r_1$ and $r_2$ the intrinsic cellular growth rates with $K_1$ and $K_2$ the carrying capacities of the sensitive and tolerant cells, respectively. Here, $\alpha_{11}$, $\alpha_{12}$, $\alpha_{21}$ and $\alpha_{22}$ refer to the payoff values associated with the effect of a sensitive cell on another sensitive cell, the effect of a tolerant cell on a sensitive cell, the effect of a sensitive cell on a tolerant cell and the effect of a tolerant cell on another tolerant cell, respectively.

**Modified Lotka-Volterra - Heterotypic cultures**:

Assuming that the cell population growth is as follows:

$$\frac{dX}{dt} = r_X X\left(1\ - \frac{X}{K_X}\right) \qquad (3)$$

where "X" represents the number of cells of the given species, $r_X$ represents the proliferation rate constant of the cells also termed as the growth rate constant, and $K_X$ , known as the carrying capacity. Assuming that the growth rate and carrying capacity are independent of seed population and time, we used all the individual population growth experiments' mean data to fit the simulated dynamics and obtained the growth rate and carrying capacity of sensitive and tolerant cells to be $r_S$ = 0.0268 $hr^{-1}$, $K_S$ = 119590 cells; $r_T$ = 0.0377 $hr^{-1}$, $K_T$ = 100520 cells (simulation fits at Fig.5A); where S represents sensitive cell population and T represents tolerant cell population. Notice that the tolerant cells have higher growth rate but smaller value of carrying capacity in comparison with sensitive cells. However, the fitted model captures the initial trends but tend to diverge significantly after 24 hrs indicating that the assumption of constant carrying capacity is not quite right. To address this we assumed a functional form for the carrying capacities which we shall discuss next.

Using these growth parameters, for the mixed co-culture dynamics of these cells we assumed the following about the nature of cell growth in a co-culture: (1) Growth rate is an inherent property of a species and is independent of the presence of another species in the culture. (2) No phenotypic switching occurs, i.e., cells maintain their identity of being sensitive or resistant throughout the experiment. (3) The carrying capacity of the sensitive and resistant subpopulations can be influenced in presence of the other.

Based on these assumptions, the dynamics of sensitive and tolerant cells during their co-culture is modelled as follows:

$$\frac{dS}{dt} = r_S S\left(1\ - (\frac{aS}{K_S{}'} + \frac{bT}{K_T'})\right) \qquad (4)$$

$$\frac{dT}{dt} = r_T T\left(1\ - (\frac{cS}{K_S{}'} + \frac{dT}{K_T'})\right) \qquad (5)$$

where $cc_S' = cc_S H(r \rightarrow s)$

$$cc_T' = cc_T H(s \rightarrow r)$$

Hills functions ($H(r \rightarrow s), H(s \rightarrow r)$) represent the change in carrying capacity of one subpopulation in presence of the other. For any 2 species X and Y, the form of a hills function for inhibition would be of the form $H(x \rightarrow y) = \frac{1}{1+x}$ , to be interpreted as carrying capacity of Y decreases as the fraction of X ($x$ ) increases in the culture. Similarly, for enhancement of carrying capacity, the functional form will be such that $H(x \rightarrow y) = \frac{x}{1+x}$. Depending on the form of the hill's function we find that in the case of competitive scenario, the best fit model indicate that the tolerant cells facilitate the growth of sensitive cells (see Supplemental section for additional details).

As mentioned in the manuscript, the dynamics of sensitive and tolerant cells during their co-culture is modelled as follows:

$$\frac{dS}{dt} = r_S S \left(1 - (\frac{aS}{K_S{'}} + \frac{bT}{K_T'})\right) \quad (6)$$

$$\frac{dT}{dt} = r_T T \left(1 - (\frac{cS}{K_S{'}} + \frac{dT}{K_T'})\right) \quad (7)$$

with $cc_S{'} = cc_S H(r \to s)$ and $cc_T' = cc_T H(s \to r)$.

Hills functions ($H(r \to s), H(s \to r)$) represent the change in carrying capacity of one subpopulation in presence of the other. For any 2 species X and Y, the form of a hills function for inhibition would be of the form $H(x \to y) = \frac{1}{1+x}$ , to be interpreted as carrying capacity of Y decreases as the fraction of X ($x$ ) increases in the culture. Similarly, for enhancement of carrying capacity, the functional form will be such that $H(x \to y) = \frac{x}{1+x}$. The four terms of the form $\frac{a*S}{cc_S{'}}$in the above equations have two components. The first part, $\frac{S}{cc_S{'}}$ , represents the amount of growth species S has already undergone relative to its carrying capacity. The constant $a$ represents the effect of the growth on the equation included in. For example, a negative value of $a$ will indicate that the growth rate of S will increase with increasing S. Similarly, a negative value of $b$ indicates that the growth of S is facilitated by the growth of R.

We considered all 4 possible combinations (competition, symbiosis, predation by sensitive cells, and predation by resistant cells) of the effect of each species on the carrying capacity of the other. One would expect positive values for the parameters (a to d) as the growth of one species expectedly depletes the resources and hence has a negative effect on the growth of the population. Given below is a table summarizing the results for each of the four cases and the corresponding estimated values of the parameters a to d, obtained by fitting the corresponding equations to 3-week co-cultured data.

| **Interaction** | $\boldsymbol{H(r \to s)}$ | $\boldsymbol{H(s \to r)}$ | $\boldsymbol{a}$ | $\boldsymbol{b}$ | $\boldsymbol{c}$ | $\boldsymbol{d}$ | **Figure** |
|---|---|---|---|---|---|---|---|
| Competition | $\frac{1}{1+r}$ | $\frac{1}{1+s}$ | 1.4661 | -0.3451 | 1.8579 | 1.0223 | 1A |
| Predation by sensitive cells | $\frac{r}{1+r}$ | $\frac{1}{1+s}$ | 0.0906 | 0.8790 | 1.6453 | 1.2356 | 1B |
| Predation by resistant cells | $\frac{1}{1+r}$ | $\frac{s}{1+s}$ | 0.7832 | 0.0048 | 0.9788 | 0.0318 | 1C |
| Symbiosis | $\frac{r}{1+r}$ | $\frac{s}{1+s}$ | 0.2707 | 0.0964 | 0.6020 | 0.1779 | 1D |

Case1 (Competition): The carrying capacity of each species is being decreased by the other in the co-culture. The negative value of b indicates that the growth rate of sensitive cells increases

with increasing fraction of resistant cells. Thus, resistant cells facilitate the growth of sensitive cells.

Case 2 (Predation by sensitive cells): The carrying capacity of sensitive cells is increased by the presence of resistant cells, in other words, higher the fraction of resistant cells, higher the saturation level of resistant cell population. In this case, all the estimated values of parameters a, b, c and d are positive, indicating that the growth rate decreases in general with increasing number of cells, which is the expected behaviour of logistic growth. Interestingly, the reduction in growth rate of sensitive cells due to the increase in their own population (a) is quite small.

Case 3 (Predation by resistant cells): In this case, the effect of resistant population on sensitive cells (b) is negligible. At the same time, the effect of sensitive cells on resistant cell growth rate (c) is positive, suggesting that the growth rate of resistant cells reduces when their carrying capacity is increased by sensitive cells.

Case 4 (Symbiosis): The carrying capacities of both species is increased by the presence of the other species. In this case, all constants are positive and of smaller magnitudes than the previous cases; the interpretation of this observation is unclear.

In all cases, c and d are always positive, suggesting that the growth rate of resistant cells is always decreased with an increase in population size. When mutual competition effect was assumed in carrying capacities, the growth rate of sensitive cells was increased by resistant cells. When predation of sensitive cells was assumed, the sensitive cell growth rate was not strongly affected by their own fraction, but was affected by the fraction of the slow growing resistant cells. In the third case, when resistant cells were assumed to be predatory, their effect on the growth rate of sensitive cells became negligible. All these cases seem to point towards a dominance of sensitive cells, with similarities to a predatory interaction, when both species are co-cultured.

**Methods:** To fit the model equations to the population data, MATLAB function *fminseach* was used, with the cost function as the norm of the difference between experimental data matrix and the corresponding simulation data matrix.

### Density game like model

We observed that the maximum population of both sensitive and tolerant cells (**Suppl. Fig. 8A-D**) varies as we change the initial seeding population. This further led us to conclude that carrying capacities, for both sensitive and tolerant cells, may not be a constant, unlike the assumptions in Zhang et al 2017. The effects of limited resources on the population growth, *in vitro*, could be captured via the growth rate (Cressman, 1990a, 1990b; Cressman and Garay, 2003) or by assuming varying carrying capacity with population density or with time (Marshall and Quental, 2016; Novak et al., 2013; Yukalov et al., 2014, 2012). In our case, varying carrying capacity model best fit the experimental data for heterotypic cultures. We do not use this model to fit monotypic cultures since it requires the existence of both phenotypes in pure cultures.

The mLV type model with varying carrying capacities we assume was of the following form:

$$\frac{dS}{dt} = r_S\, S\left(1 - \frac{S}{K_S}\right) \#(8)$$

$$\frac{dT}{dt} = r_T\, T\left(1 - \frac{T}{K_T}\right) \#(9)$$

where $K_S = \left(A_{SS}\frac{S}{S+T} + A_{ST}\frac{T}{S+T}\right)e^{-C_S t}$ and $K_T = \left(A_{TS}\frac{S}{S+T} + A_{TT}\frac{T}{S+T}\right)e^{-C_T t}$ are the carrying capacities for sensitive and tolerant cells, respectively, and $A_{SS}, A_{ST}$, $A_{TS}$, $A_{TT}, C_S$ and $C_T$ are parameter constants. The above forms for carrying capacities is similar to Novak et al., (2013) in which payoff values using an alternative evolutionary game theory (EGT) perspective influence the carrying capacities of respective populations. Novak et.al. assume that for the model which can be written as

$$\frac{dx_i}{dt} = r_i x_i\left(1 - \frac{x_T}{k_i}\right) \#(10)$$

$x_i$ represents different phenotype populations, $x_T = \sum_i x_i$ represents total population, $r_i$ is the growth rate and $k_i$ denotes carrying capacity for ith phenotype. Assuming that

$$k_i = \sum_j \alpha_{ij}\frac{x_j}{x_T}, \#(11)$$

They propose that $\alpha_{ij}$ are the payoff values that determine the competition between different phenotype populations.

It should be noted that our model differs from Novak et al. model in two aspects. i) Novak et al. model uses $x_T$ term in numerator (see Eqns. 10 and 11) as opposed to $x_i$ term that we have used in our model (see Eqns. 8 and 9 ). We assume that this does not affect our interpretation of payoff values. ii) Carrying capacities in our model depend on time because of time dependent exponential terms. The payoff values in our model can be written as follows:

$$\alpha_{ij} = A_{ij}e^{-C_i t} \#(12)$$

Where $i = j = \{S, T\}$. This implies that the payoff values in our model changes with time.

*Heterotypic cultures:*

**Suppl. Fig. 7A** and **7B** show the fitting for sensitive and tolerant cell data, respectively. Looking at payoff values from **Suppl. Fig. 7G**, our model indicates that the presence of tolerant phenotype has a positive effect on sensitive one whereas sensitive phenotype effect on tolerant is adverse. Also, the positive effect of cells on similar phenotype cells ($\alpha_{SS}$ and $\alpha_{TT}$) is an order of magnitude higher and hence more significant than effect of cells on different phenotype cells ($\alpha_{ST}$ and $\alpha_{TS}$).

**Suppl. Fig. 7D** and **7E** show the model fit obtained for mixed culture data in the presence of cisplatin. Some important conclusions that can be drawn from the corresponding payoff values (**Suppl. Fig. 7H)** are: i) Inter-phenotypic effects ($\alpha_{ST}$ and $\alpha_{TS}$) are negligible when compared with intra-phenotypic effects ($\alpha_{SS}$ and $\alpha_{TT}$). For example, initial value of $\alpha_{SS}$ is approximately

13 times higher than $\alpha_{ST}$. iii) The positive effect of tolerant on tolerant phenotype ($\alpha_{SS}$) is almost 30 times stronger than the effect of sensitive on sensitive phenotype ($\alpha_{TT}$). Thus, in the presence of cisplatin, tolerant phenotype is the dominant player.

It is interesting to compare the relative decrease in growth rates for mixed-phenotype culture when cisplatin dose is introduced., We observe that decrease in tolerant cells' growth rate is relatively sharper than that of sensitive (see Table 1 below). We propose that the sharper decrease in growth rate is the cost tolerant phenotype pay for developing better resistance for cisplatin. This better resistance can be the result of better cooperation between tolerant-tolerant cells (compared to sensitive-sensitive cells), which is manifested as significantly higher payoff value ($\text{i.e. } \alpha_{TT} \gg \alpha_{SS}$).

The parameter values that fit the mixed culture of sensitive and tolerant cells under various proportions after 3 weeks in the absence and presence of Cisplatin are given in Table 1.

*Table 1: Model Parameters for density game like model*

| **Culture** | $r_1$ | $r_2$ | $A_{11}$ | $A_{12}$ | $A_{21}$ | $A_{22}$ | $C_1$ | $C_2$ |
|---|---|---|---|---|---|---|---|---|
| Mixed Co-culture without | 0.0253 | 0.028 | $3.025 \times 10^5$ | $3.22 \times 10^4$ | $-1.01 \times 10^4$ | $5.17 \times 10^5$ | 0.015 | 0.020 |
| Mixed Co-culture with Cisplatin | 0.0197 | 0.016 | $1.98 \times 10^4$ | $1.638 \times 10^4$ | 0.0 | $6.45 \times 10^5$ | 0.0107 | 0.037 |

*Phenotype-switch Model with stress response*

$$\frac{K_a}{K_b} = K_0(1 + aC_{Str}) \quad (13)$$

where $K_0$ is the ratio between $K_a$ and $K_b$ in absence of stress and *a* represents the sensitivity of phenotype switching to stress variation. Furthermore, $K_b$ is assumed to be constant. Both, $K_{GS}$ and $K_{GT}$ decrease linearly with stress according to equations 9 and 10,

$$K_{GS} = K_{GS_0}(1 - bC_{Str}) \quad (14)$$

$$K_{GT} = K_{GS_0}(1 - gC_{Str}) \quad (15)$$

where $K_{GS_0}$ and $K_{GT_0}$ are growth rate constants in absence of stress and *b*, *g* represent the sensitivity of the growth rate constants to stress variation.

*Fitting of the phenotype-switch model to experimental growth trends*

The fitting was performed using the global optimization protocol, Genetic Algorithm (GA) [6], as implemented in the package GA in R [7]. First, a local optimization using the Nelder-

Mead nonlinear optimization was performed for each growth trend individually. The fitted parameters that showed reasonable agreement with experiments were used to estimate the range of variation for each parameter, and were provided as lower and upper bounds to the GA routine. In GA, the population size was set to 50 and the maximum number of iterations was 5000. All other settings were kept at their default values. The optimization was considered to have converged once the best solution did not change for more than 100 steps. The fitting was performed separately to the growth data from the sensitive and tolerant cells alone, as well as the mixed cultures, leading to separate parameters for each of these experiments. The fitting error was estimated as the mean of the absolute deviations between the observed and the fitted trends, normalized by the cell population range.

**Fig. 5B** and **5C** and **Suppl. Fig. 8** show the agreement between the fitted and observed growth trends for different types of systems (pure sensitive or tolerant cells, mixed cultures where counting begun immediately or after three weeks of co-culture). For each system, a single set of parameters fitted the different growth curves (various seeding populations or mixing proportions) with reasonable agreement (please see the fitting errors in **Suppl. Fig. 8C, 8F** and **8I**). Among the sensitive or tolerant only systems, the worst error was observed for the lowest seeding population (**Suppl. Fig. 8C**, 1250 cells/well). It is possible that at low seeding populations, stochastic effects would dominate, and the systems may be best modeled using stochastic differential equations instead of deterministic ones. In the mixed system where cell counting was performed immediately, the error increased proportionally with the amount of tolerant cells in the system, indicating that the tolerant cell behavior may be more complex than what the model describes. The errors were relatively low for the systems after three weeks of co-culture, since the majority of the tolerant cells were predicted to have switched their phenotypes to the sensitives after three weeks, as will be discussed later. Hence, the growth dynamics after three weeks was dominated by the sensitive phenotype, even when seeded with high population of green fluorescent cells that were descended from tolerant ancestors.

*Effects of phenotype switching and stress give rise to diverse game-theoretical strategies in mixed cell populations*

We have analyzed the cellular behavior using two types of theoretical models based on different perspectives. The game theoretic models give the broad strategies such as competition and cooperation adopted by the cells for survival and maximizing growth. The phenotype-switch model analyzes the cell behavior at a finer level by giving insights into the phenotypic plasticity and environmental factors that determine growth. We next asked whether these two models can be integrated; in other words, can the phenotype-switch model explain the game-theoretical strategies displayed by the cellular phenotypes?

Due to the complexity of the model equations, it proved impossible to analytically derive the game theoretical equations from the phenotype-switch model. We therefore derived the LV and density games model parameters by numerically fitting the growth rates from the phenotype-switch model to the game theory equations for different time-windows (**Suppl. Figs. 10 and 11A-C)**. Both for small (12 hours) and very large time-windows (6 days), the fitting seemed worse, while moderate length time windows such as 4-5 days (**Suppl. Figs. 10 and 11B**), the fitting appeared reasonable between the two models. Also, the mLV model parameters derived from the 4 day time window are in agreement with those obtained from fitting the density-game like model to the experimental data, as shown in **Suppl. Fig. 11E**. To explore the changing landscape of cooperative strategies adopted by the cells, we plotted the LV parameters $\alpha_{12}$ and $\alpha_{21}$ as function of fitting time window and sensitive to tolerant seeding ratio (**Fig. 5J** and **5K**). As seen before, both small and large fitting time windows show high fitting error (>0.08). This suggests that at small timescales (< 24 hours), the cellular behavior as captured by the Phenotype-switch model deviates from the LV model whereas at larger timescales ranging (1-5 days), the two models were in agreement. For very large timescales (> 6 days), the two models began to diverge since a constant set of game-theory parameters may not describe the growth behavior over such a long time period. The fitting error is also higher when the proportion of tolerant cells increase in the system suggesting that the tolerant cells may introduce more complexity to the growth behavior than the combination of the two models could capture.
