## Supplemental Figures for "Suppressing chemoresistance in lung cancer via dynamic phenotypic switching and intermittent therapy"

Supplementary Fig. 1

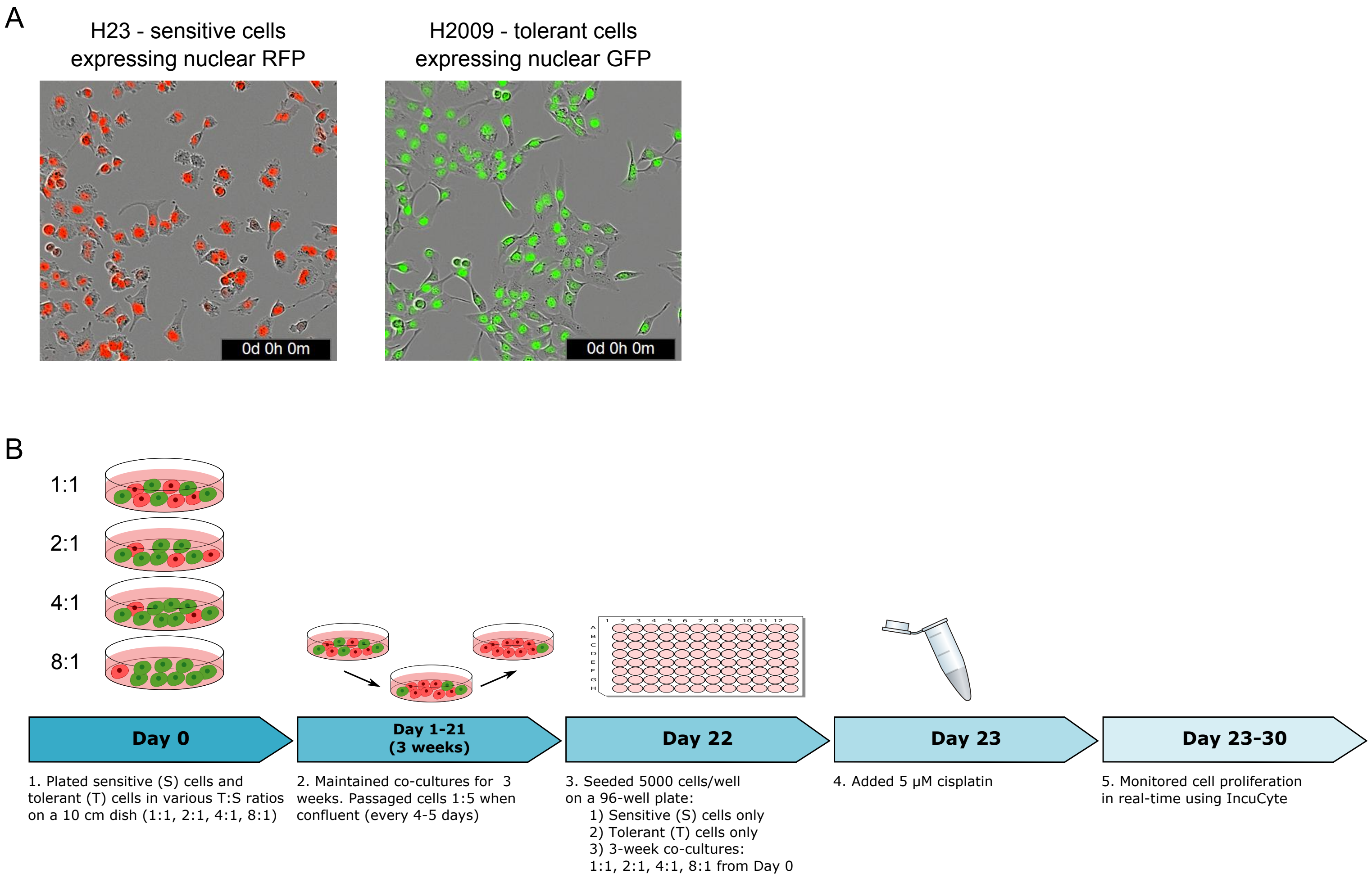

Supplementary Fig. 2

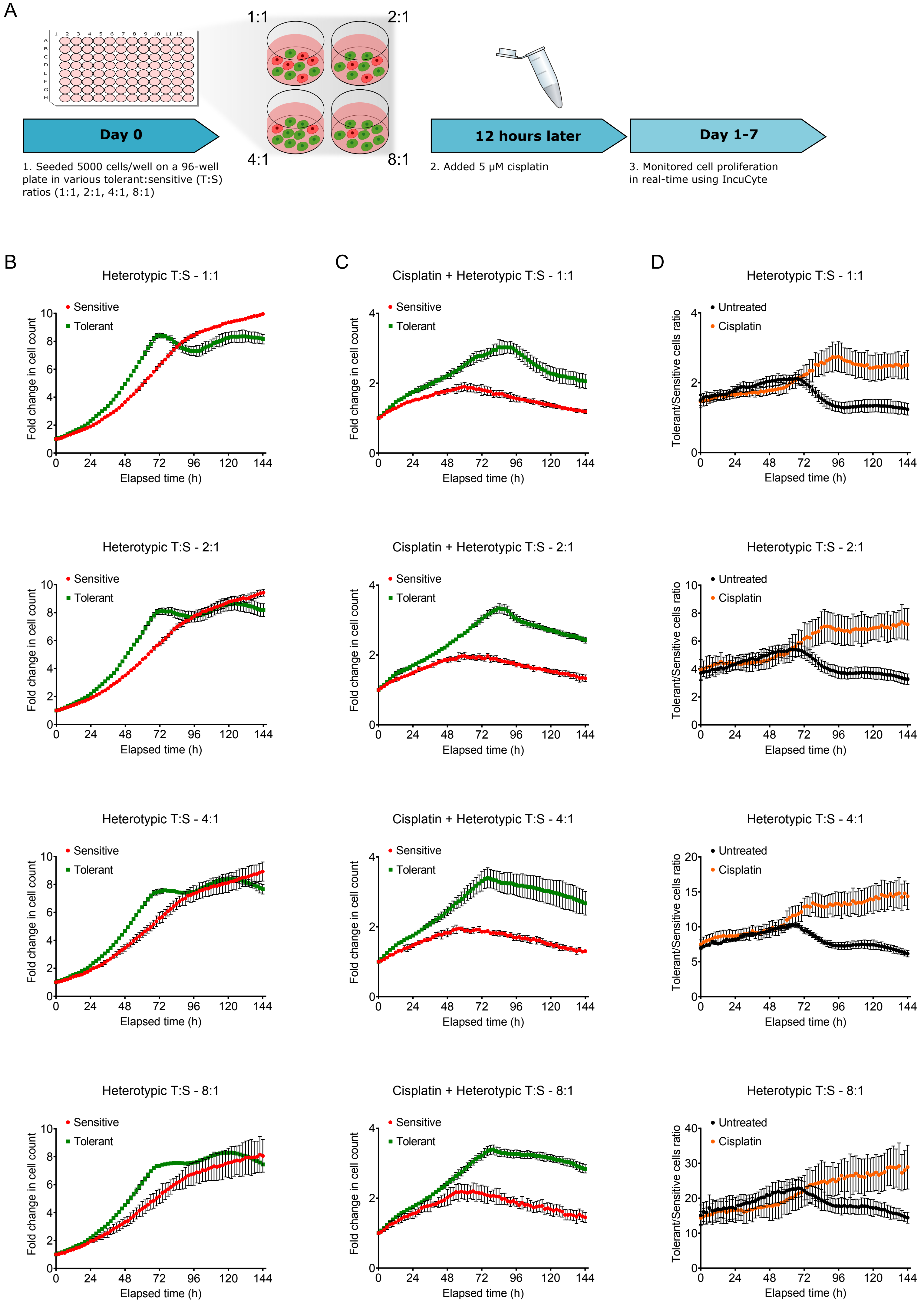

Supplementary Fig. 3

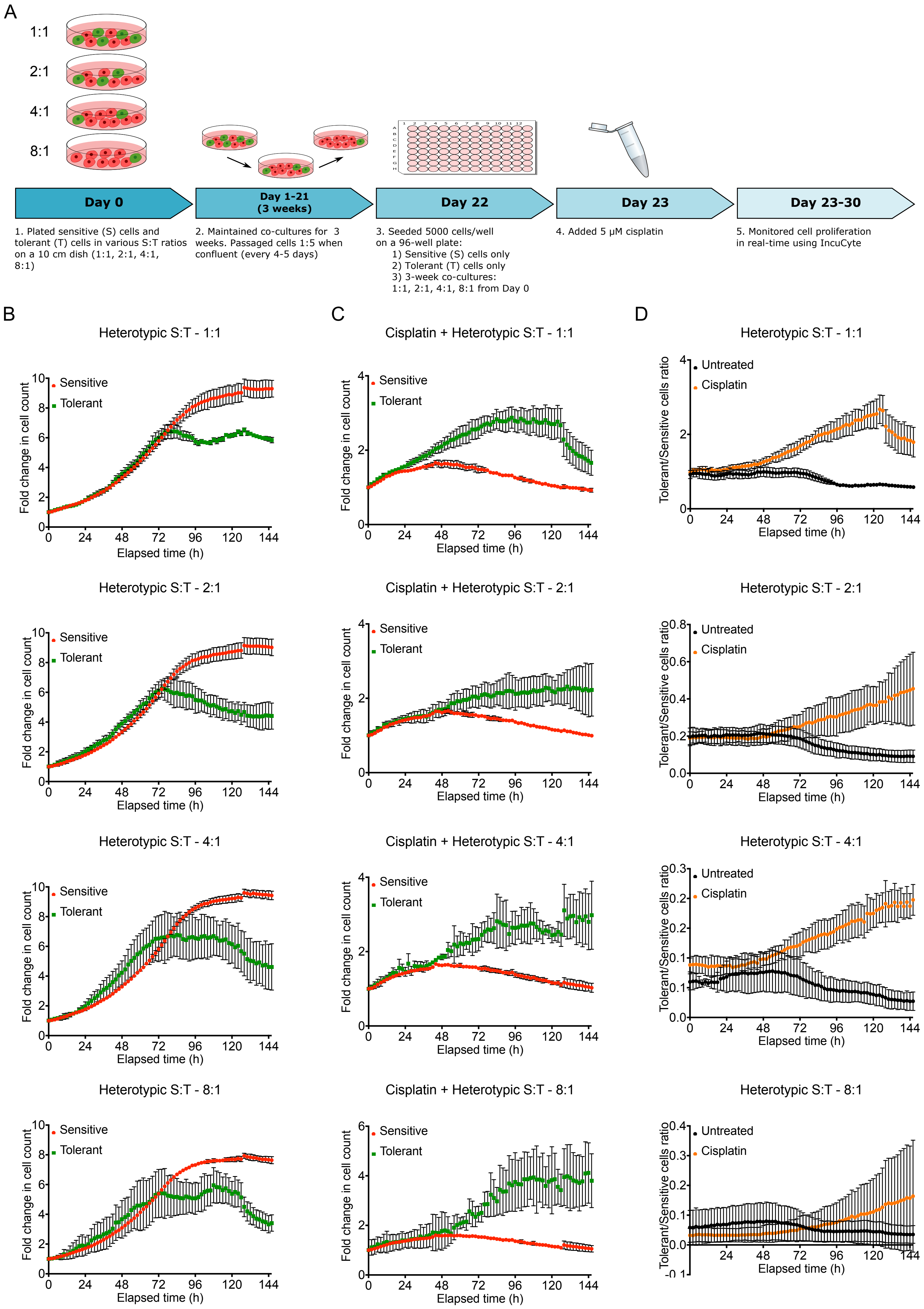

Supplementary Fig. 4

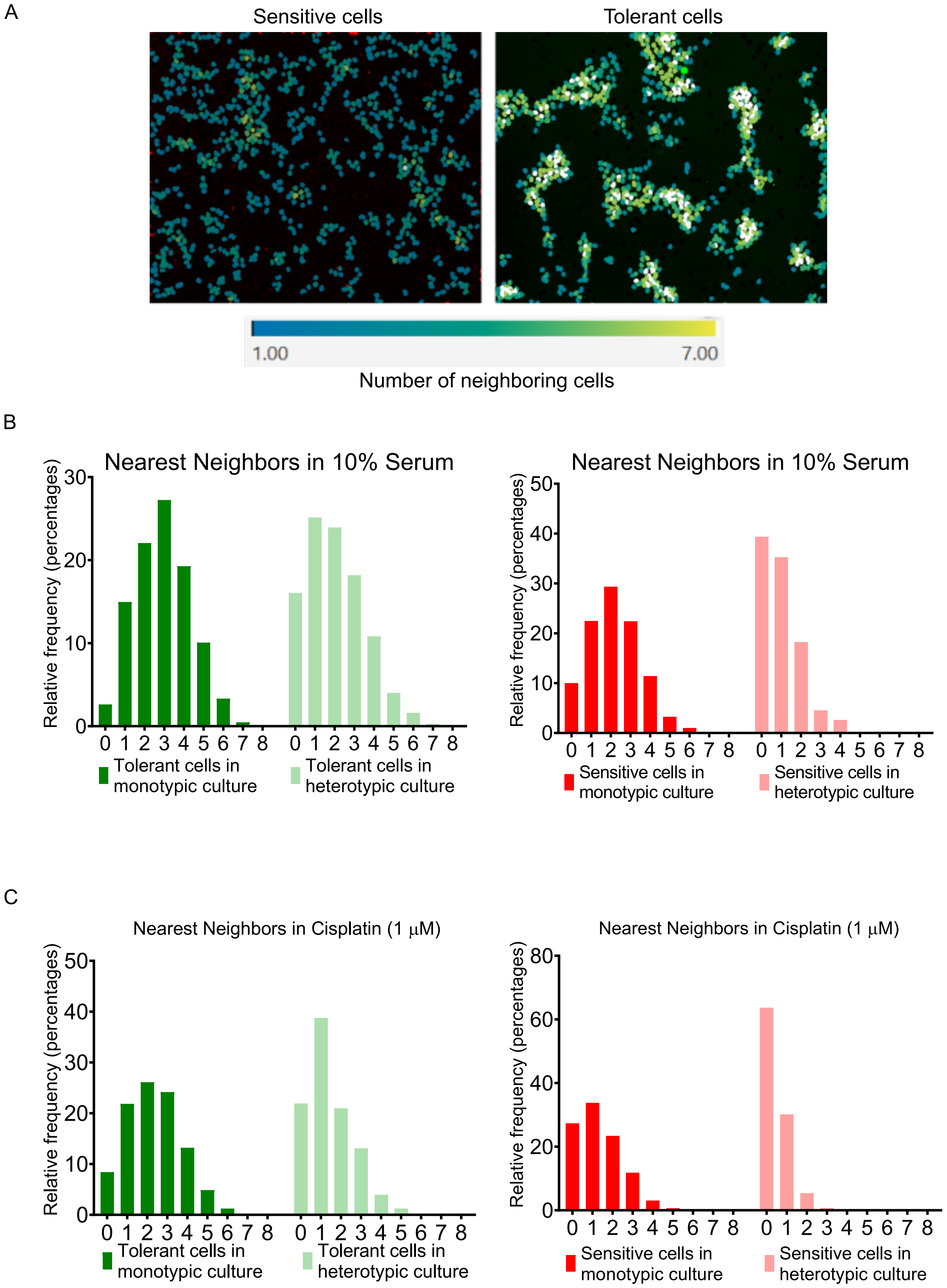

Supplementary Fig. 5

A

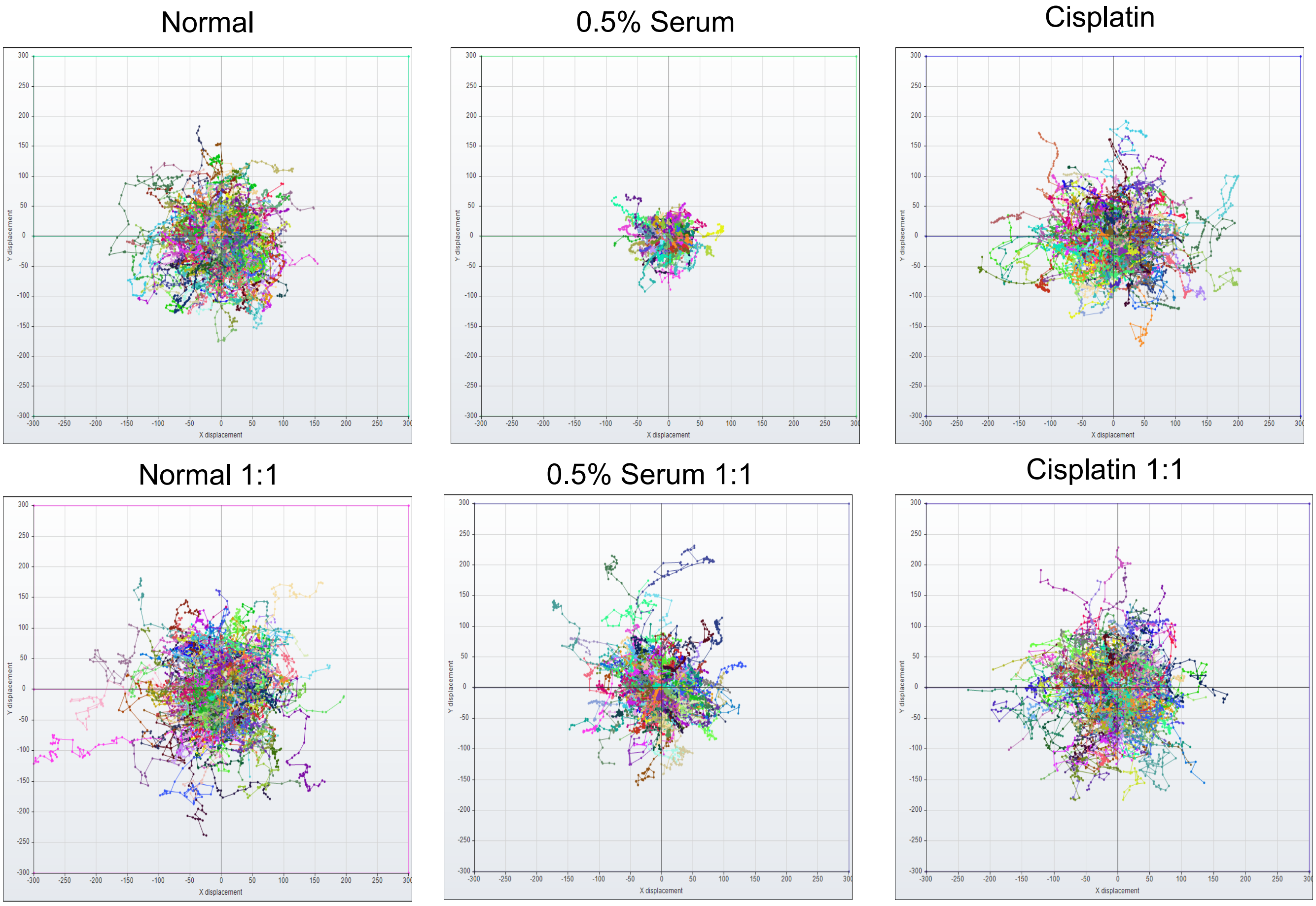

B

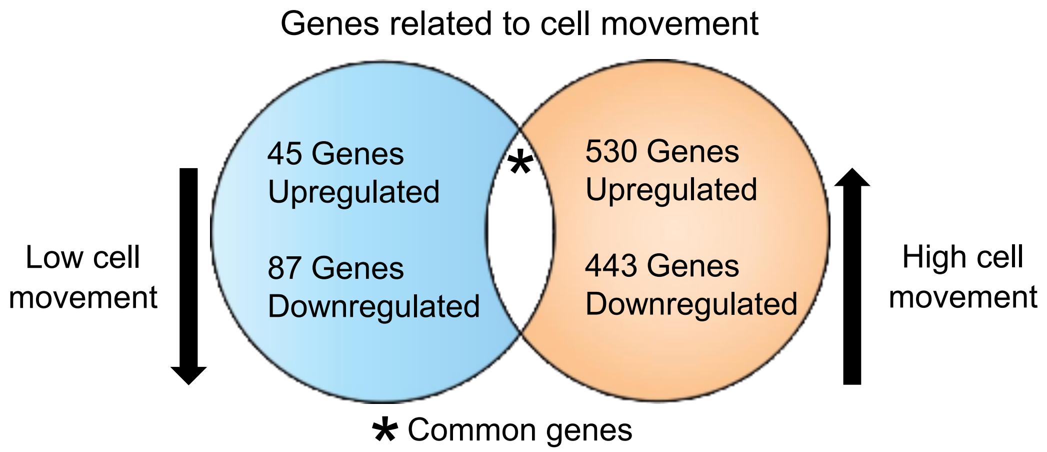

C

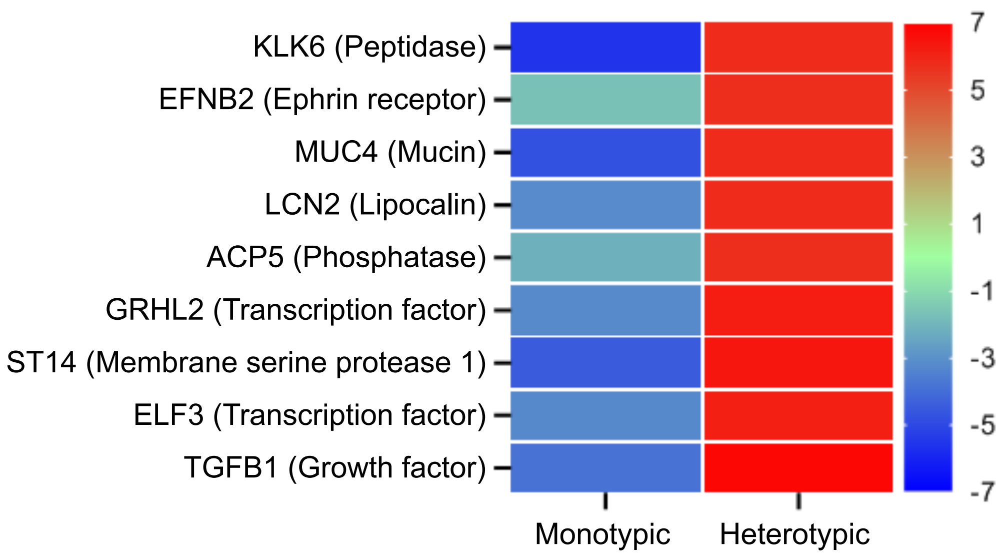

Supplementary Fig. 6

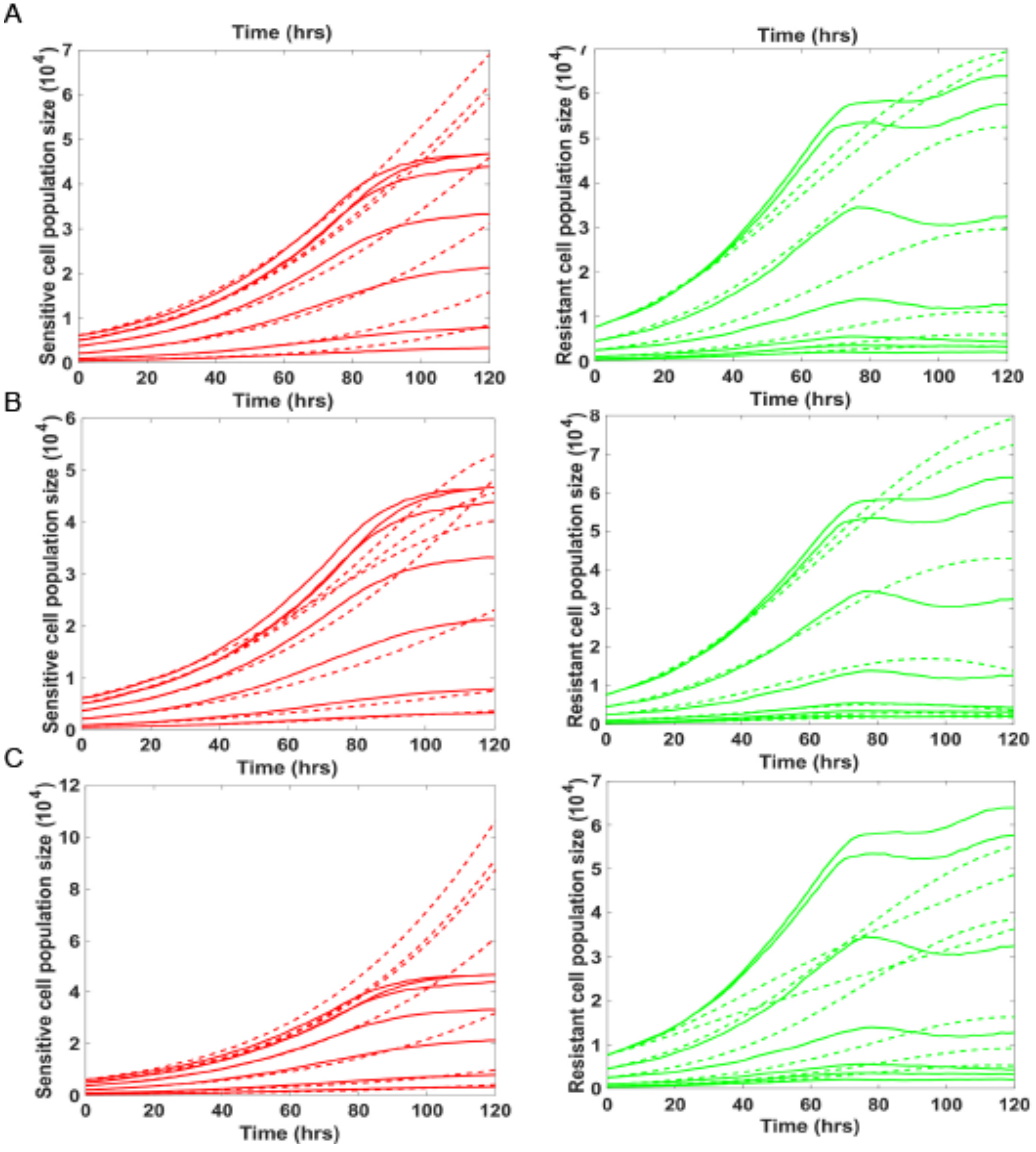

Supplementary Fig. 7

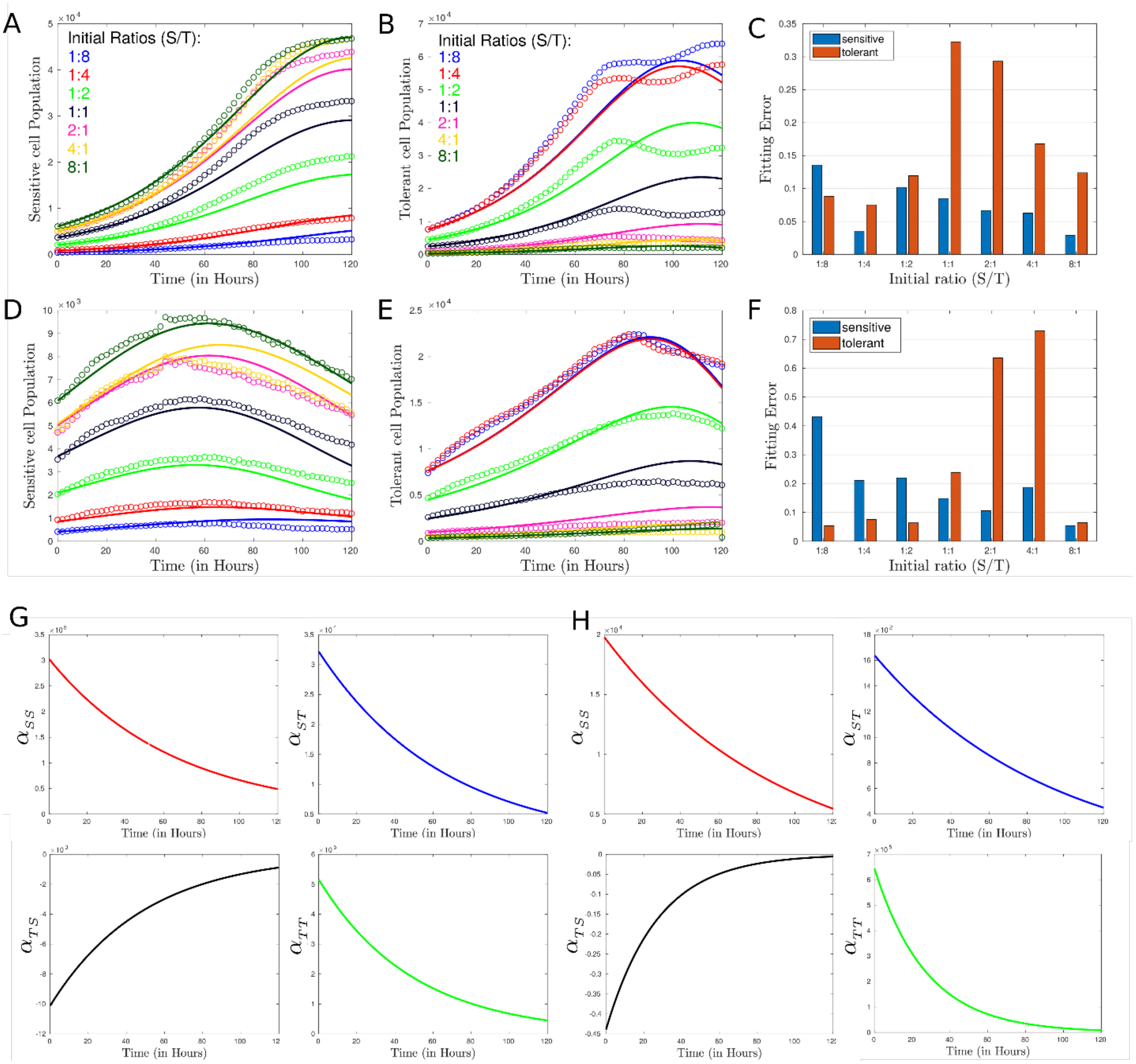

### Supplementary Fig. 8

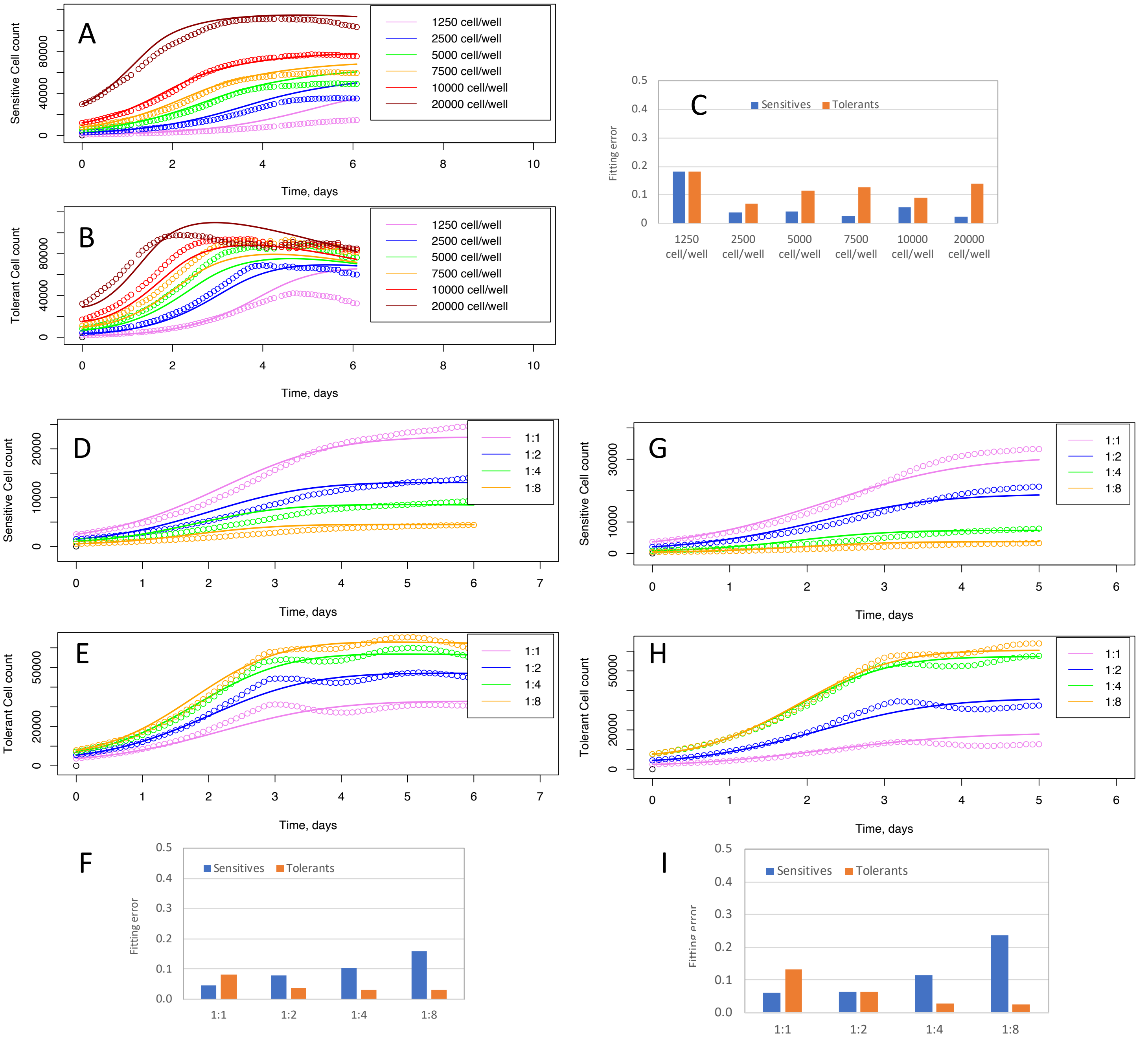

J

| Condition | $K_0$ | $K_b$ | $K_{Gs0}$ | $K_{Gt0}$ | $K_s$ | $K_{sd}$ | a | b | g |
|---|---|---|---|---|---|---|---|---|---|
| Sensitives only | 9.51E-03 | 1.01 | 0.626 | 0.093 | 2.94E-04 | 6.06E-03 | 0.050 | 0.036 | 0.026 |
| Tolerants only | 1.00E-02 | 1.98 | 1.012 | 0.108 | 2.56E-04 | 2.70E-04 | 0.044 | 0.036 | 0.043 |
| Mixed | 4.87E-02 | 3.54 | 0.713 | 0.682 | 7.14E-04 | 5.67E-03 | 0.045 | 0.037 | 0.015 |
| Mixed 3 weeks | 4.98E-02 | 2.66 | 0.695 | 0.313 | 6.72E-04 | 5.38E-03 | 0.050 | 0.038 | 0.016 |

Supplementary Fig. 9

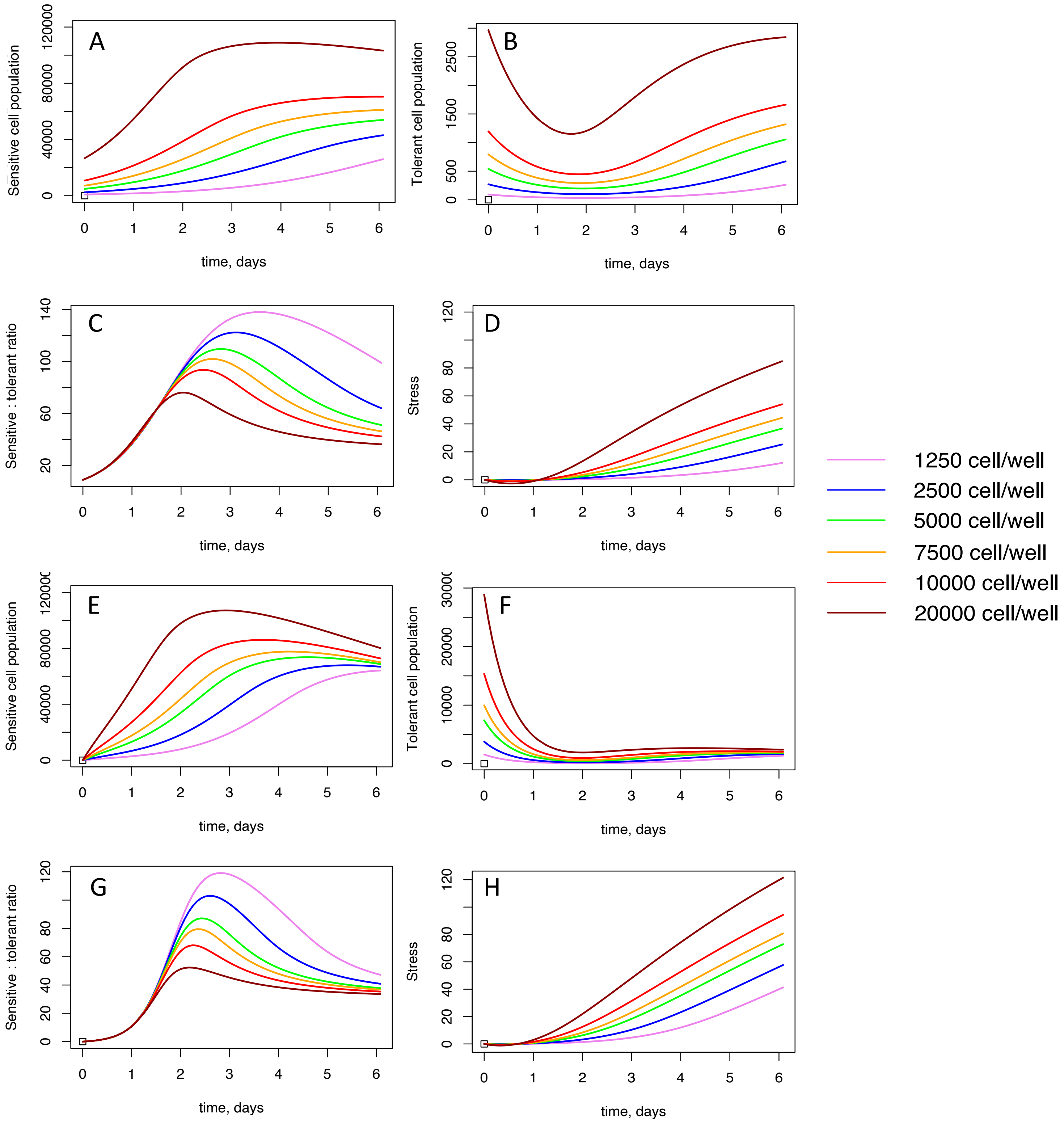

### Supplementary Fig. 10

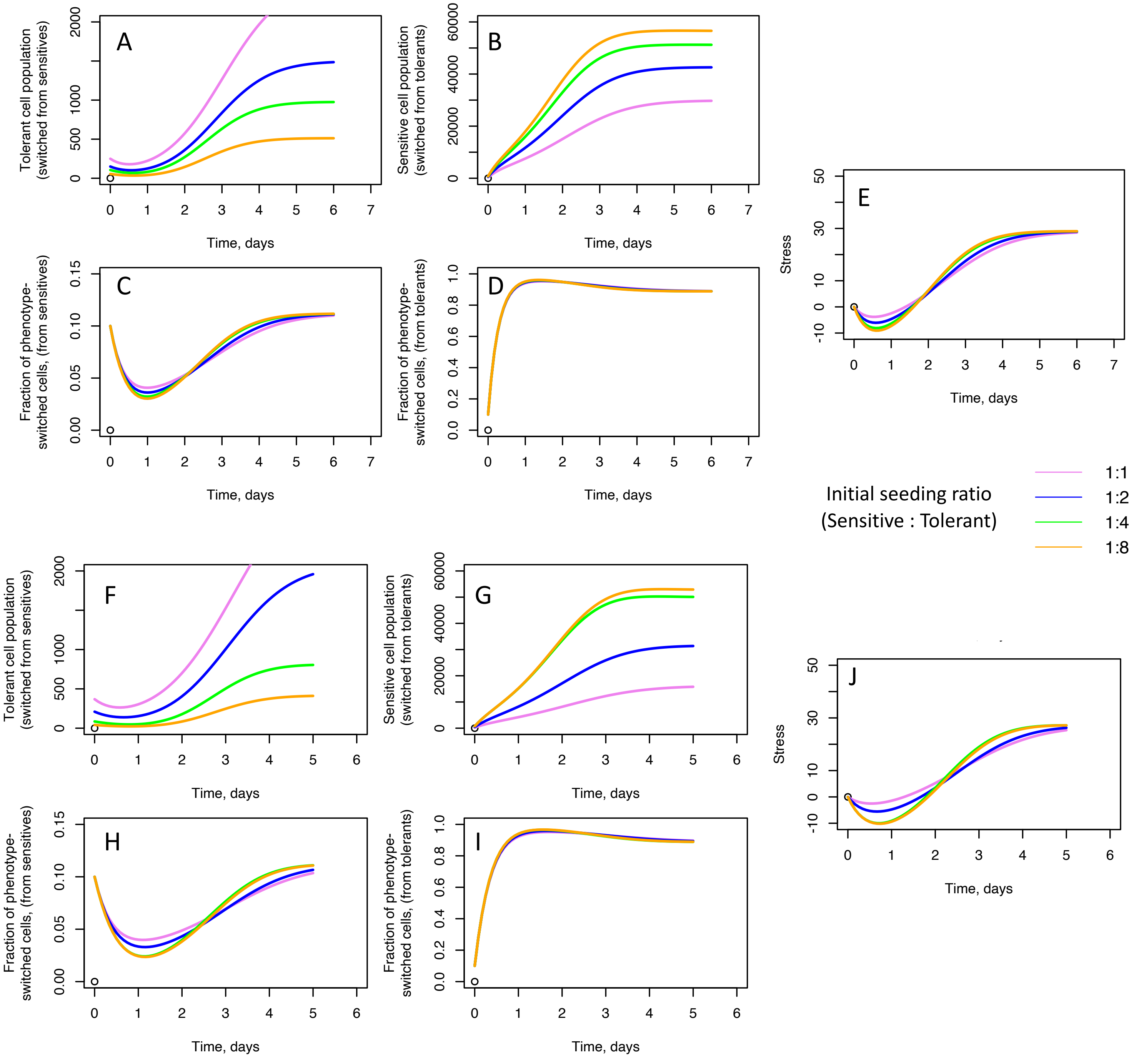

### Supplementary Fig. 11

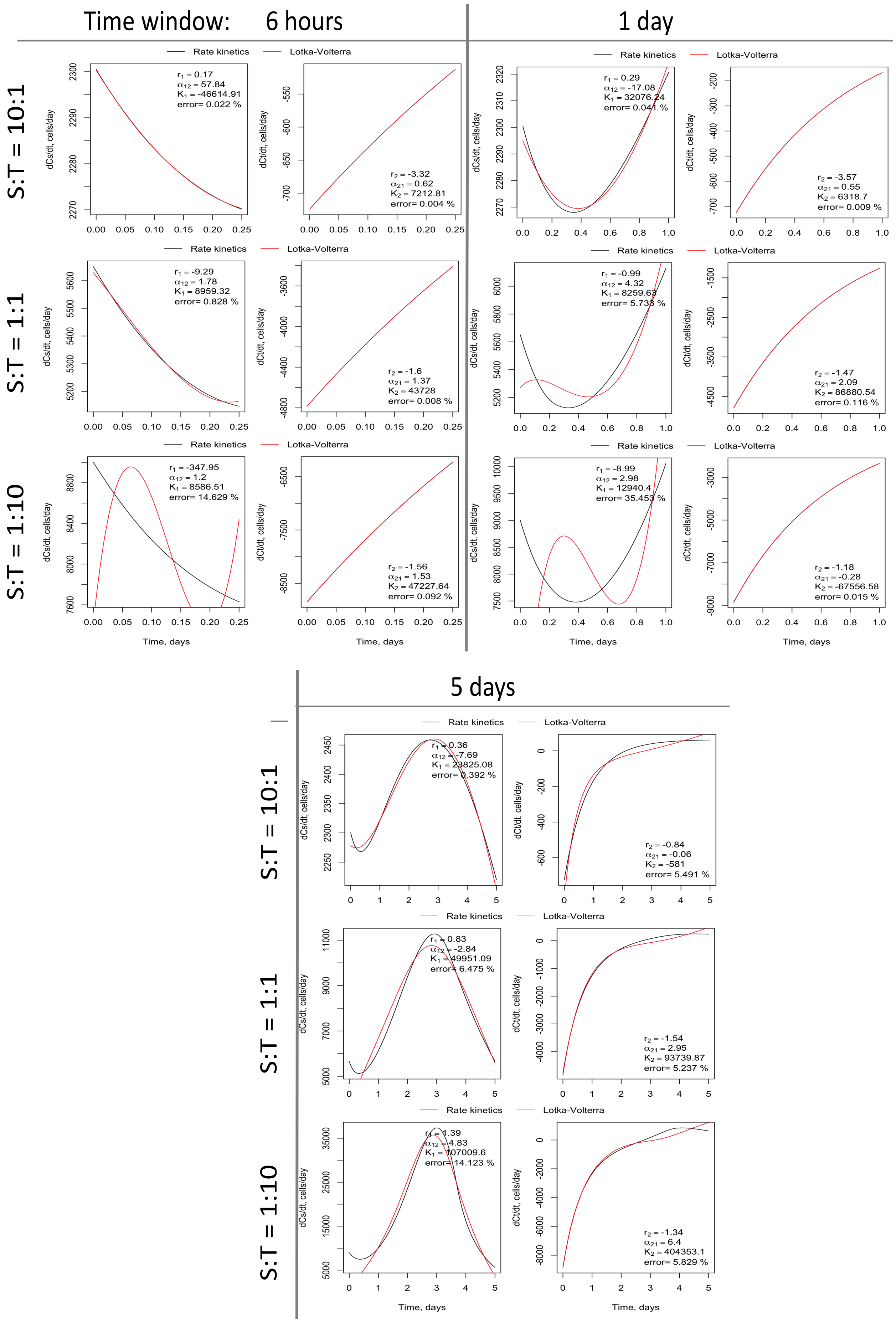

### Supplementary Fig. 12

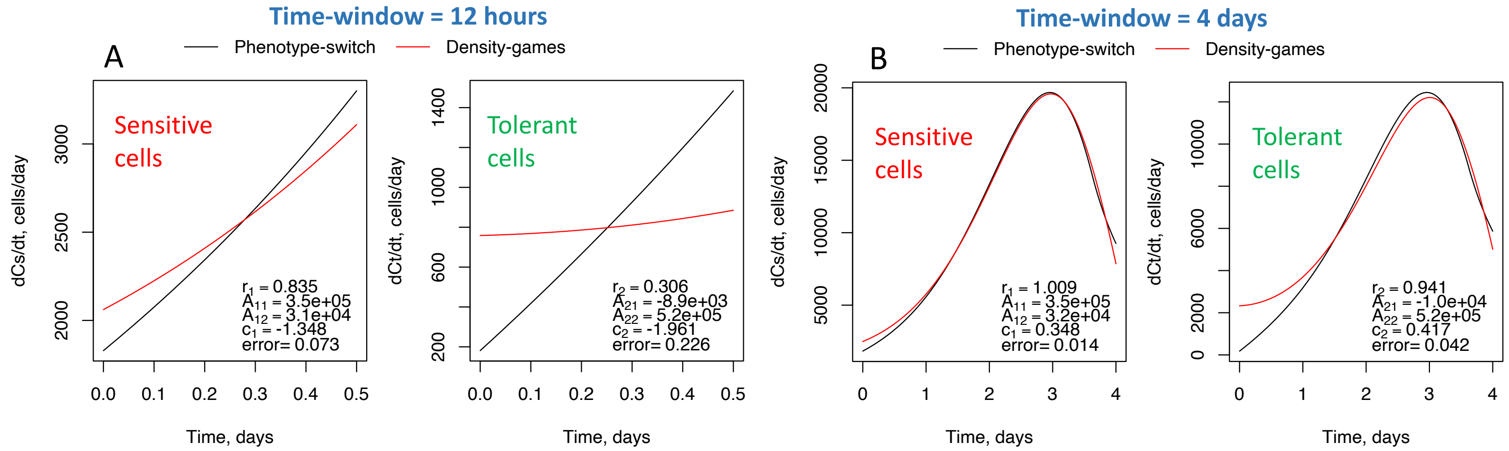

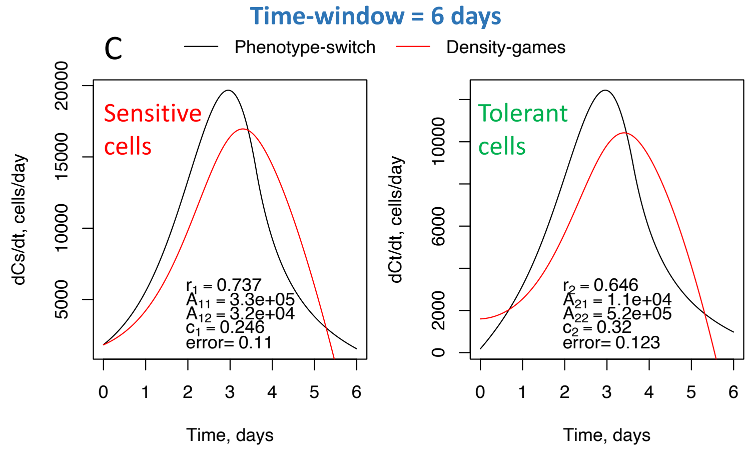

D

**Density-games model**

*($x_1$, $x_2$: sensitive & tolerant cell population)*

$$\frac{dx_i}{dt} = r_i x_i \left(1 - \frac{x_i}{k_i}\right)$$

$$k_i = \left(b_{ii}\frac{x_i}{x_1 + x_2} + b_{ij}\frac{x_j}{x_1 + x_2}\right) e^{-c_i t}$$

E

| Source | $r_1$ | $r_2$ | $A_{11}$ | $A_{12}$ | $A_{21}$ | $A_{22}$ | $C_1$ | $C_2$ |
|---|---|---|---|---|---|---|---|---|
| Fitting to exp. data | 0.607 | 0.672 | $3.025\times10^5$ | $3.22\times10^4$ | $-1.01\times10^4$ | $5.17\times10^5$ | 0.36 | 0.48 |
| Derived from Phenotype -switch model | 1.009 | 0.941 | $3.5\times10^5$ | $3.2\times10^4$ | $-1.0\times10^4$ | $5.2\times10^5$ | 0.348 | 0.417 |

Supplementary Table 1

| Row | Incubation period prior to live cell counting | Seeding ratio (Tolerant : Sensitive) | Ratio at first time point of live cell counting |
|---|---|---|---|
| 1 | 12 hours | 1:1 | ~2:1 |
| 2 | 12 hours | 2:1 | ~4:1 |
| 3 | 12 hours | 4:1 | ~7:1 |
| 4 | 12 hours | 8:1 | ~13:1 |
| 5 | 3 weeks | 1:1 | ~1:5 |
| 6 | 3 weeks | 2:1 | ~2:1 |
| 7 | 3 weeks | 4:1 | ~9:1 |
| 8 | 3 weeks | 8:1 | ~15:1 |
| 9 | 3 weeks | 1:1 | ~8:10 |
| 10 | 3 weeks | 1:2 | ~2:10 |
| 11 | 3 weeks | 1:4 | ~5:100 |
| 12 | 3 weeks | 1:8 | ~5:100 |

Supplementary Table 2. Statistical significance for Fig. 1B-D

| Figure 1B | | | | |
|---|---|---|---|---|
| T:S ratio | Time point | Statistical test | P-value | Significance |
| 1:1 | 144 h | 2-way ANOVA | <0.0001 | Significant |
| 2:1 | 144 h | 2-way ANOVA | <0.0001 | Significant |
| 4:1 | 144 h | 2-way ANOVA | <0.0001 | Significant |
| 8:1 | 144 h | 2-way ANOVA | <0.006 | Significant |

| Figure 1C | | | | |
|---|---|---|---|---|
| T:S ratio | Time point | Statistical test | P-value | Significance |
| 1:1 | 144 h | 2-way ANOVA | 0.5015 | Not significant |
| 2:1 | 144 h | 2-way ANOVA | <0.0001 | Significant |
| 4:1 | 144 h | 2-way ANOVA | <0.0001 | Significant |
| 8:1 | 144 h | 2-way ANOVA | <0.006 | Significant |

| Figure 1D | | | | |
|---|---|---|---|---|
| T:S ratio | Time point | Statistical test | P-value | Significance |
| 1:1 | 144 h | 2-way ANOVA | <0.0001 | Significant |
| 2:1 | 144 h | 2-way ANOVA | <0.0001 | Significant |
| 4:1 | 144 h | 2-way ANOVA | <0.0001 | Significant |
| 8:1 | 144 h | 2-way ANOVA | <0.0001 | Significant |

Supplementary Table 3. Statistical significance for Fig. 2A

| Figure 2A | | | | |
|---|---|---|---|---|
| Experiment condition | Time point | Statistical test | P-value | Significance |
| Tolerant-conditioned media | 72 h | 2-way ANOVA | 0.019 | * |
| Sensitive-conditioned media | 72 h | 2-way ANOVA | 0.001 | *** |

Supplementary Table 4. Statistical significance for Fig. 4A-C

| Figure 4A-C | | | |
|---|---|---|---|
| S:T ratio | Statistical test | P-value | Significance |
| 1:1 | Wilcoxon two-tailed | <0.0001 | Significant |
| 2:1 | Wilcoxon two-tailed | <0.0001 | Significant |
| 4:1 | Wilcoxon two-tailed | <0.0001 | Significant |

Supplementary Table 5. Statistical significance for Supplementary Fig. 2B-D

| Supp. Figure 2B | | | | |
|---|---|---|---|---|
| T:S ratio | Time point | Statistical test | P-value | Significance |
| 1:1 | 144 h | 2-way ANOVA | <0.0001 | Significant |
| 2:1 | 144 h | 2-way ANOVA | <0.0001 | Significant |
| 4:1 | 144 h | 2-way ANOVA | <0.0001 | Significant |
| 8:1 | 144 h | 2-way ANOVA | 0.1569 | Not significant |

| Supp. Figure 2C | | | | |
|---|---|---|---|---|
| T:S ratio | Time point | Statistical test | P-value | Significance |
| 1:1 | 144 h | 2-way ANOVA | <0.0001 | Significant |
| 2:1 | 144 h | 2-way ANOVA | <0.0001 | Significant |
| 4:1 | 144 h | 2-way ANOVA | <0.0001 | Significant |
| 8:1 | 144 h | 2-way ANOVA | <0.006 | Significant |

| Supp. Figure 2D | | | | |
|---|---|---|---|---|
| T:S ratio | Time point | Statistical test | P-value | Significance |
| 1:1 | 144 h | 2-way ANOVA | <0.0001 | Significant |
| 2:1 | 144 h | 2-way ANOVA | <0.0001 | Significant |
| 4:1 | 144 h | 2-way ANOVA | <0.0001 | Significant |
| 8:1 | 144 h | 2-way ANOVA | <0.0001 | Significant |

Supplementary Table 6. Statistical significance for Supplementary Fig. 3B-D

| Supp. Figure 3B | | | | |
|---|---|---|---|---|
| S:T ratio | Time point | Statistical test | P-value | Significance |
| 1:1 | 144 h | 2-way ANOVA | <0.0001 | Significant |
| 2:1 | 144 h | 2-way ANOVA | <0.0001 | Significant |
| 4:1 | 144 h | 2-way ANOVA | <0.0001 | Significant |
| 8:1 | 144 h | 2-way ANOVA | 0.1569 | Not significant |

| Supp. Figure 3C | | | | |
|---|---|---|---|---|
| S:T ratio | Time point | Statistical test | P-value | Significance |
| 1:1 | 144 h | 2-way ANOVA | <0.0001 | Significant |
| 2:1 | 144 h | 2-way ANOVA | <0.0001 | Significant |
| 4:1 | 144 h | 2-way ANOVA | <0.0001 | Significant |
| 8:1 | 144 h | 2-way ANOVA | <0.006 | Significant |

| Supp. Figure 3D | | | | |
|---|---|---|---|---|
| S:T ratio | Time point | Statistical test | P-value | Significance |
| 1:1 | 144 h | 2-way ANOVA | <0.0001 | Significant |
| 2:1 | 144 h | 2-way ANOVA | <0.0001 | Significant |
| 4:1 | 144 h | 2-way ANOVA | <0.0001 | Significant |
| 8:1 | 144 h | 2-way ANOVA | 0.3784 | Not significant |
